# Refphase: Multi-sample reference phasing reveals haplotype-specific copy number heterogeneity

**DOI:** 10.1101/2022.10.13.511885

**Authors:** Thomas BK Watkins, Emma C Colliver, Mathew R Huska, Tom L Kaufmann, Emilia L Lim, Kerstin Haase, Peter Van Loo, Charles Swanton, Nicholas McGranahan, Roland F Schwarz

## Abstract

Most computational methods that infer somatic copy number alterations (SCNAs) from bulk sequencing of DNA analyse tumour samples individually. However, the sequencing of multiple tumour samples from a patient’s disease is an increasingly common practice. We introduce Refphase, an algorithm that leverages this multi-sampling approach to infer haplotype-specific copy numbers through multi-sample reference phasing. We demonstrate Refphase’s ability to infer haplotype-specific SCNAs and characterise their intra-tumour heterogeneity, to uncover previously undetected allelic imbalance in low purity samples, and to identify parallel evolution in the context of whole genome doubling in a pan-cancer cohort of 336 samples from 99 tumours.

## Background

As a consequence of genomic instability, cancers accumulate somatic mutations [1]. These include somatic copy number alterations (SCNAs) affecting focal genomic segments, chromosome arms, or entire chromosomes, and whole genome doubling (WGD) events that alter the entire karyotype. SCNAs change the number of physical copies of a given genomic region and often result in aneuploidy, which affects up to 90% of solid tumours [2,3]. These mutational events may be clonal, shared by all cancer cells, or subclonal and thus present only in a subset of cells, resulting in intra-tumour heterogeneity (ITH) [4,5]. It is now clear that single-sample sequencing studies derived from single biopsies are often insufficient to capture the extent of mutational heterogeneity and the field increasingly relies on multi-sample bulk and single-cell analyses from the same tumour to better describe this complexity. Such studies have permitted the mapping of the landscape of clonal and subclonal SCNAs [6–12], and have revealed a relationship between SCNA intra-tumour heterogeneity and poor prognosis in multiple tumour types [6,8,13–16]. Additionally, chromosomal instability and aneuploidy have been associated with and proposed as causes of cancer drug resistance [17] and linked to metastasis [18–21]. Inference of SCNAs and quantification of their intra-tumour heterogeneity is therefore of clinical importance and a prerequisite for understanding tumour evolution.

SCNAs are typically identified in an allele-specific manner from DNA sequencing or single nucleotide polymorphisms (SNP) array data using two measures: the log read-depth ratio (LogR) of a genomic locus between the tumour and a matched normal sample, which informs total copy number estimates; and the B-allele frequency (BAF) at heterozygous SNPs, which informs allelic imbalance (AI) estimates [22]. BAF and LogR profiles are sensitive to sequencing noise and sample purity. The difficulty of characterising SCNAs in low tumour purity bulk samples can lead to many being discarded [23]. Imposing such purity thresholds on analyses is likely problematic as sample purity co-segregates with other important clinical covariates and survival [23]. Therefore, approaches that accurately resolve copy number states in low purity samples are of clinical interest.

Additionally, while BAF and LogR allow inference of allele-specific SCNAs, the resulting copy number profiles and underlying SNPs are often not phased, meaning that SCNAs cannot be assigned to the physical haplotypes they reside on. Instead, allele-specific copy number states are typically reported in an unphased major/minor configuration, where major and minor refer to the greater and lesser copy number at a genomic locus respectively. However, phasing of SCNAs can offer additional insights into the clonality of mutational events, giving resolution on whether the same parental allele has been gained or lost across different samples in a single tumour.

Statistical phasing utilises large collections of genotypes [24] and local linkage disequilibrium structure to phase SNPs [25,26]. Multiple groups have previously implemented statistical phasing approaches in the context of whole-genome sequencing (WGS) [27], single-sample bulk sequencing studies [27,28] and in single-cell studies, using DNA [29] and RNA [30]. However, while highly accurate locally, statistical phasing accuracy rapidly decreases with increasing genomic distance, limiting the genomic span within which a SNP and the corresponding SCNA can accurately be assigned to its haplotype-of-origin. Therefore, statistical phasing for SCNA detection is mostly restricted to WGS data, ignoring a large proportion of cancer genomics studies based on whole-exome (WES) or targeted panel sequencing. Additionally, while single-cell sequencing removes the complications of sample purity, it is so far not routinely used in clinical studies and genomic coverage typically remains low, impeding allele-specific copy number readouts.

To address these challenges, we present Refphase. Refphase implements multi-sample reference phasing [9] to infer haplotype-specific copy number states, rescue previously undetected SCNAs in low purity samples, and standardise quantification of SCNA-based ITH. Refphase is, to our knowledge, the first long-range phasing algorithm applicable to all of multi-sample WGS, SNP array, exome and targeted sequencing data. Unlike statistical phasing, Refphase does not require reference haplotype panels or large collections of genotypes. Instead, it leverages the common germline background between multiple samples from the same patient to phase heterozygous SNPs and SCNAs. We have previously used reference phasing to describe MSAI, independent SCNAs that occur on opposite haplotypes in different samples from the same tumour, and to identify parallel and convergent copy number events [8,9].

Refphase integrates SCNAs pre-segmented using single-sample approaches [22,31–33] across multiple samples from the same tumour. It takes input from ASCAT [22] and other common copy number callers and provides output compatible with MEDICC2 [34], enabling streamlined processing of multi-sample copy number data from raw read counts to event-based SCNA phylogenies.

Here, we demonstrate Refphase’s ability to infer SCNAs and characterise their intra-tumour heterogeneity; implement its grouping functionality to compare the SCNA landscapes in primary and metastatic tumour samples from a single patient’s disease; showcase its ability to uncover previously undetected AI in low purity samples; and use it to identify parallel evolution in the context of whole genome doubling in a pan-cancer cohort.

## Results

### Refphase algorithm

Refphase takes as input a copy number segmentation for each of ***N*** tumour samples from the same patient which it processes in four discrete steps or modules (Figure 1, Methods).

**Figure 1:**
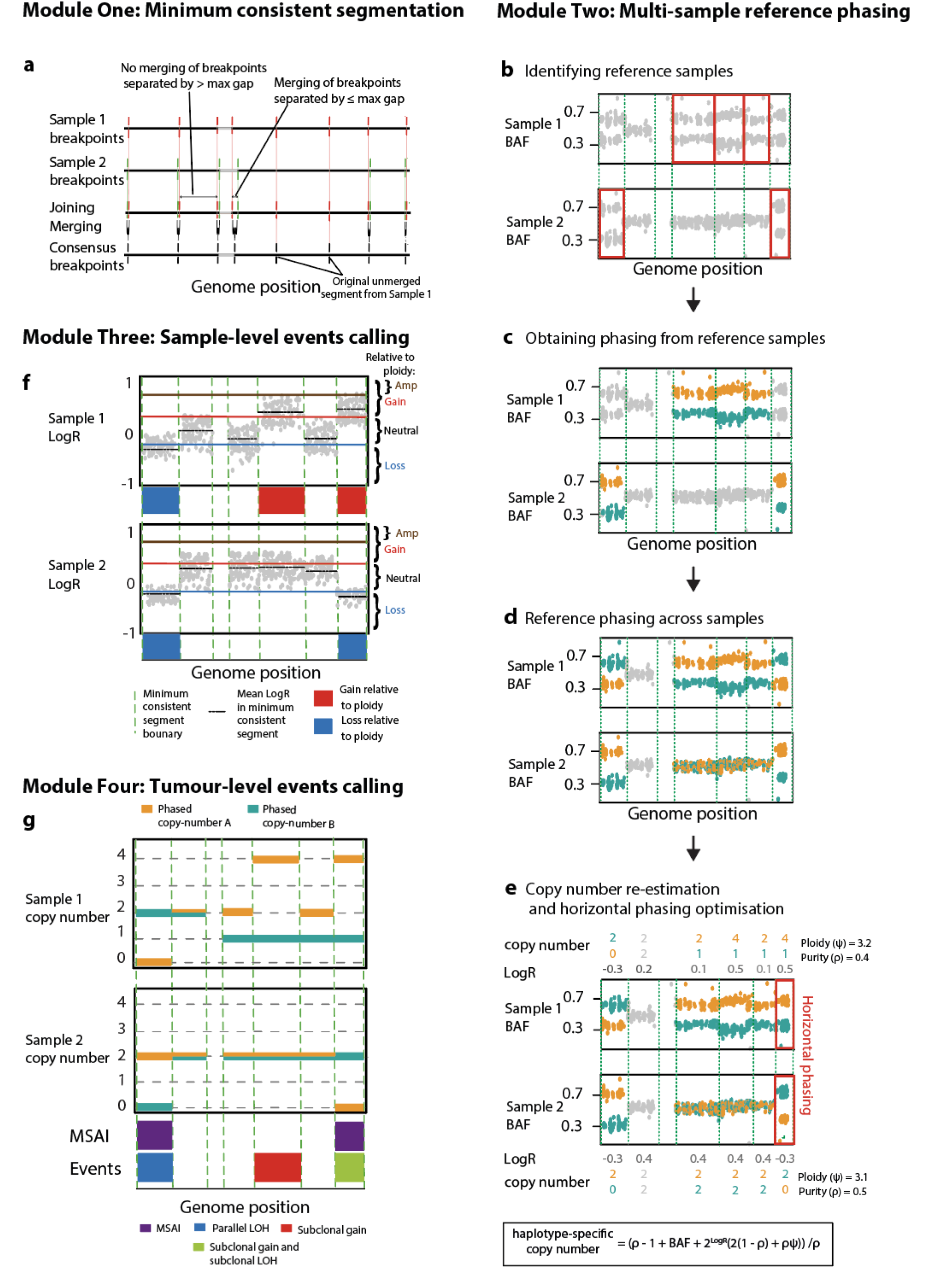
Overview of Refphase algorithm. **a)** Refphase creates a minimum consistent segmentation across the single-sample segmentations input for each tumour. **b)** In each segment in which at least one sample had allelic imbalance in the tumour input, an optimal reference sample for phasing is determined. **c)** The phasing of each reference sample is derived from its BAF. **d)** Phasing is then applied to the BAFs in all other samples which are not the reference. **e)** Allele-specific copy numbers are re-estimated for each sample utilising the reference phasing, and the most parsimonious phasing solution along each chromosome is then chosen in horizontal phasing optimization. **f)** In each segment, event categories relative to the input ploidy of the corresponding sample are defined using LogR values. **g)** Tumour-level events are called and intra-tumour heterogeneity metrics calculated.

First, a minimum consistent segmentation is created from the union of all start and end positions of the input copy number segmentations of each tumour sample. Breakpoints with distance smaller than a user-defined maximum gap size (default = 100kbp) are subsequently merged. This step excludes breakpoints that belong to the same sample-of-origin to preserve focal gains and losses present in the original samples (Figure 1a, Methods) and yields a final set of **m** bins.

Next, using this set of **m** bins, multi-sample reference phasing is performed (Figures 1b-e). Heterozygous SNPs are either defined by the user or identified from the BAF values of the normal sample (Methods). For each bin **m_i_,** the sample with the highest degree of AI is then identified, which will act as a reference ***n*_ref_** for **m_i_** (Figure 1b, Methods). This reference sample ***n*_ref_** is then used to assign alleles of all heterozygous SNPs within **m_i_** to the “A” or “B” haplotype based on their BAFs being greater than or less than a threshold of 0.5 (Figure 1c). This haplotype assignment in turn is then applied to all other tumour samples for the same bin **m_i_** (Figure 1d) and each tumour sample is assessed for the presence of any additional previously undetected AI using an effect size threshold based on Cohen’s d (Methods). Haplotype-specific integer copy number states are then re-estimated for all reference phased segments (Figure 1e, Methods).

Reference phasing as described above accurately phases the haplotypes of all samples for each bin relative to one another, but independently of other bins. To phase bins along the genome “horizontally”, we next estimate this horizontal phasing by using an evolutionary criterion. Briefly, the assignment of heterozygous SNPs to “A” and “B” haplotypes for all bins within a single chromosome is chosen to minimise the number of copy number breakpoints across all tumour samples [34,35] (Figure 1e, Methods, Supplementary Figure 1). The relative phasing between samples from the previous step therefore remains unchanged.

The third module then uses the re-estimated copy number and updated AI states to determine whether a segment from a tumour sample may be categorised as an SCNA relative to the ploidy of that specific tumour sample, in keeping with both previous genomics studies [8,9,36] and clinical practice [37]. This relative-to-ploidy classification enables the comparison of the same area of the genome in bin **m_i_** between samples with differing ploidies and classifies segments as either an amplification, gain, neutral, or loss relative to sample ploidy (Figure 1f, Methods). Loss of heterozygosity (LOH), copy-neutral loss of heterozygosity (CNLOH) and homozygous deletion events are also classified (Methods).

The fourth and final module integrates the multi-sample reference phasing and relative-to-ploidy classification to produce a tumour-level estimate of SCNA event clonality across all samples and to infer the presence of mirrored subclonal allelic imbalance (MSAI) (Figure 1g). All events are then summarised as clonal (present in all tumour samples without MSAI) or subclonal (present in only a subset of tumour samples, or in all samples but with MSAI detected) (Methods).

### Refphase identifies mirrored subclonal allelic imbalance and parallel evolution

We first demonstrated the functionality of Refphase by analysing WES data from three spatially separated primary tumour samples and a matched normal from a non-small cell lung cancer patient CRUK0034 from the TRACERx 100 cohort [8]. All three tumour samples were pre-segmented with ASCAT [22] and subjected to reference phasing. Dividing the genome into 178 bins, Refphase identified either clonal or subclonal AI in 88.6% of the genome, with the remaining 11.4% of the genome allelically balanced. Figure 2 shows parts of the Refphase output for this tumour. A more complete output example is available in Supplementary Figure 2. For this example and the remainder of the text, we refer to unphased copy number states in major/minor configuration as e.g. 4/1 and to phased copy number states as e.g. 1|4, in line with genotype notation conventions.

**Figure 2:**
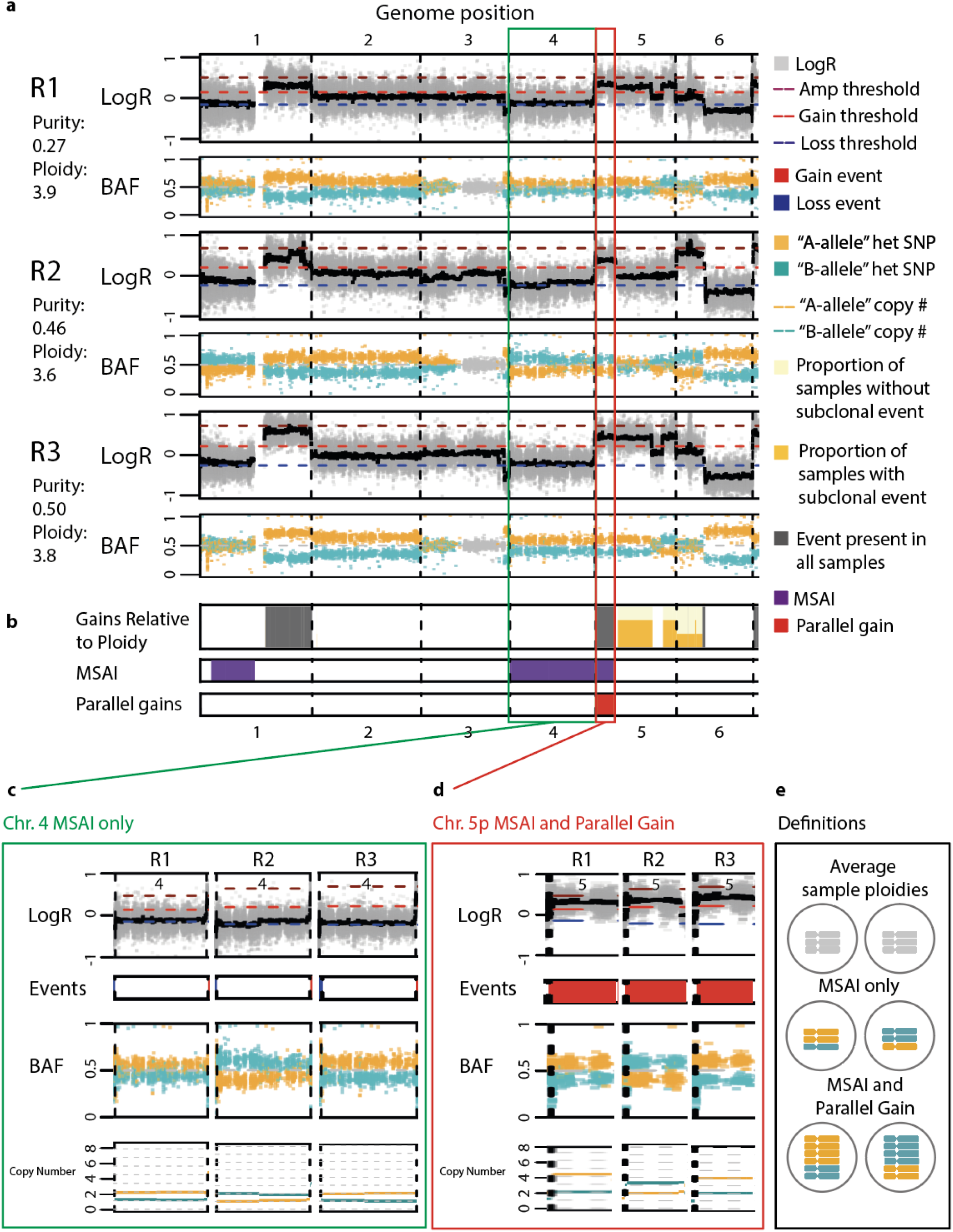
Detection of mirrored subclonal allelic imbalance and parallel evolution. **a)** LogR and BAF tracks in chromosomes 1 to 6 from tumour CRUK0034. LogR tracks show LogR values in light grey points. The black line shows the median LogR within a minimum consistent segment. BAF tracks show phased BAF as either orange “A” haplotype points or blue “B” haplotype points. Unphased BAF values are shown as light grey points. **b)** SCNA summary tracks showing (top) gains relative to ploidy. The full height grey bar indicates that a gain is identified in every sample from tumour CRUK0034. A light yellow background indicates the presence of a subclonal gain, and the height of the stacked darker yellow bar indicates the proportion of samples in which a subclonal gain is present. (middle) Track indicates the presence of MSAI between at least two samples, shown by purple fill. (bottom) Track indicates the presence of parallel gains, shown by red fill. **c)** MSAI detected from tumour CRUK0034 affecting chromosome 4. **d)** Parallel evolution of chromosome arm 5p gain from tumour CRUK0034. **e)** Schematic of the copy number states related to MSAI and parallel evolution.

While some SCNAs (e.g. gain on 1q) are clonal and present in all samples from the tumour, Refphase also identifies substantial inter-sample heterogeneity (Figure 2a-b, Supplementary Figure 2). For example, 3p demonstrates AI only in R2, where even after reference phasing, no AI was detectable in the other samples, as can be seen from the well mixed BAF values for both haplotypes (orange and blue) on 3p in R1 and R3. Refphase identifies subclonal SCNA events including subclonal gain of 5q affecting R1 and R3, and a subclonal gain of 6p affecting R2.

Reference phasing also permitted the identification of both MSAI and parallel evolution in this tumour. One additional copy of chromosome 4 relative to diploid (Figure 2c) (copy number state 2/1) is present in all tumour samples. Reference phasing revealed that this additional copy was derived from the “A” haplotype (orange) in samples R1 and R3, and from the “B” haplotype (blue) in sample R2, an example of MSAI [8]. A second instance of MSAI on chromosome 5p co-occurs with relative-to-ploidy gains in all three samples with copy number states of 4|2 in R1 and R3 but of 2|3 in R2 (Figure 2d). A schematic outlining the distinction between the classifications applied to the MSAI affecting chromosome 4 and the parallel gains affecting 5p can be seen in Figure 2e. In addition to the detection of MSAI on chromosome 4 and parallel evolution of 5p gain, Refphase also identified an additional instance of MSAI affecting 1p (Figure 2b).

Since MSAI and parallel events are not detectable from the analysis of single tumour samples nor from unphased data, Refphase’s ability to identify haplotype-specific copy number states and its event classifications provide a refined view of the evolution of this tumour.

### Refphase quantifies SCNA heterogeneity in complex multi-sample cases

The increasing availability of DNA sequencing data from multiple tumour samples from the same patient has begun to address complex questions regarding SCNA intra-tumour heterogeneity and changes in SCNAs during metastatic dissemination. However, there are no standardised frameworks in which such questions may be addressed and compared between studies. Refphase not only supports the quantification of SCNA clonality across all samples from the same patient but also allows standardised analysis and comparisons of user-defined subgroups of samples within a single patient’s disease. In addition, Refphase produces correctly formatted input for the state-of-the-art SCNA phylogenetic reconstruction algorithm MEDICC2 [34], as well as the option to output its own naive haplotype-specific clustering of minimum consistent segments.

Patient CRUK0063, previously analysed in work by Abbosh et al. in 2017 [38], was examined through the PEACE post-mortem study 24 hours after death. WES data from five post-mortem tumour samples (paravertebral and lung metastases) and five primary tumour samples were pre-segmented using ASCAT and subjected to reference phasing using Refphase. We leveraged Refphase’s group analysis capability to specifically investigate differences between primary and metastatic samples (Figure 3). MEDICC2 [34] was run on the Refphase output and grouped the metastasis samples together in one clade and the primary samples in a separate clade (Figure 3a). Refphase identified SCNA events present in both primary and metastatic samples (Figure 3b), present in primary samples only (Figure 3c), and present in metastatic samples only (Figure 3d).

**Figure 3:**
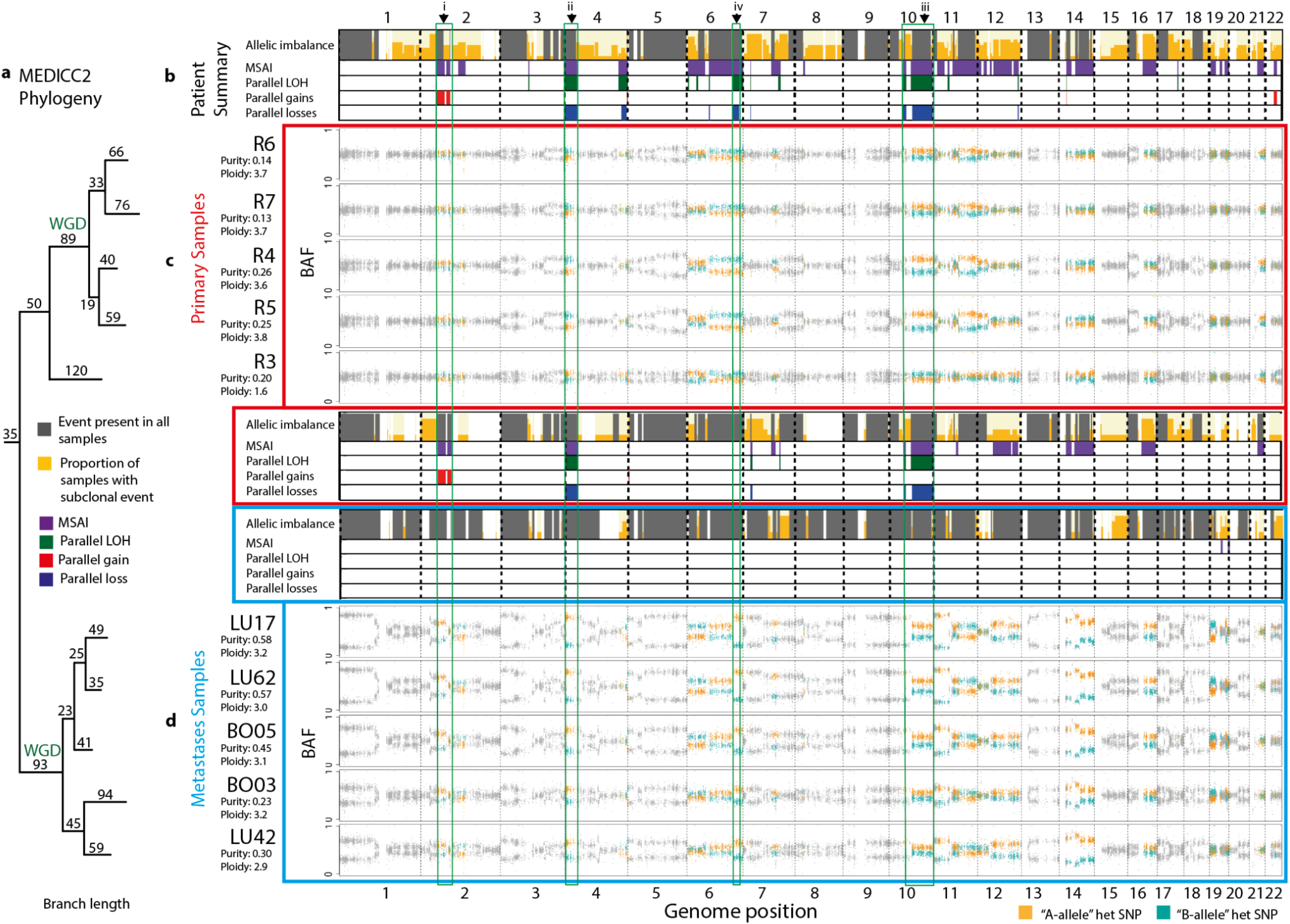
Grouping analysis of CRUK0063 multi-sample and multi-time point NSCLC case. **a)** MEDICC2 phylogeny. Multi-sample reference phased allele-specific copy number output from Refphase can be passed directly to MEDICC2 to produce a phylogenetic reconstruction. **b)** SCNA summary tracks for all samples - primary and metastases - from patient CRUK0063. **c)** BAF profiles and SCNA summary tracks for the primary samples from CRUK0063 are indicated by a red border. **d)** SCNA summary tracks and BAF profiles from five post-mortem metastatic samples with a blue border. Sample BAF tracks are ordered by their position in the MEDICC2 phylogenetic reconstruction. Green boxes, arrowheads and associated Roman numerals highlight selected examples of MSAI on chromosomes 2p, 6q, 4p and 10q, described in the main text. WGD indicates whole genome doubling events inferred by MEDICC2.

Examining all CRUK0063 samples together revealed clonal SCNAs (SCNAs here encompassing relative-to-ploidy gains and losses, and LOH events) affecting 25% of the genome, subclonal SCNAs affecting 69% of the genome, MSAI affecting 26% of the genome and parallel evolution evident in 8% of the genome. Refphase’s group analysis feature allows us to analyse the five primary and five metastatic samples separately, while using the phasing derived from all samples. In this grouped analysis, differences between the two groups emerged and enabled us to distinguish between SCNAs that are clonal and subclonal in the primary samples (primary-clonal/primary-subclonal) and those that are clonal and subclonal in the metastatic samples (metastasis-clonal/metastasis-subclonal). Metastasis-clonal SCNAs were far more prevalent than primary-clonal SCNAs, both in terms of proportion of the genome affected (55% *vs.* 36%), and of proportion of segments with any SCNA event (relative-to-ploidy gain or loss, or LOH) for which a clonal SCNA was identified, quantified within the respective group of samples (55% *vs.* 33%). Metastasis-clonal AI and LOH were particularly prevalent, affecting 72% and 34% of the genome respectively, while primary-clonal AI and LOH affected only 39% and 27% of the genome. The majority of LOH was found to be shared between primary and metastatic samples.

In line with the high levels of metastasis-clonal SCNAs in CRUK0063, the metastatic samples are also characterised by a relative absence of both MSAI and parallel evolution compared to the primary samples (proportion of genome: MSAI, 0.3% *vs.* 13%; parallel events, encompassing parallel gains, parallel losses and parallel LOH events, 0% *vs.* 5%). Examples of specific events observed when analysing the primary samples alone are visible in Figure 3 and include parallel gain of 2p (i), and multiple parallel LOH events including on 4p (ii) and 10q (iii). Specifically, primary tumour sample R3 had a different major haplotype to other primary samples at these loci. The divergence of sample R3 from other primary samples is reflected in it branching earlier than the remaining four primary samples in the accompanying MEDICC2 phylogeny. In the Abbosh et al. study [38], mutational phylogenetic analysis suggested that all metastatic samples arose from a single ancestral subclone; similar results are observed using the MEDICC2 phylogenies with all metastatic samples belonging to a single clade. There is also MSAI affecting 13% of the genome and an instance of parallel evolution of LOH affecting 6q between the primary and metastatic samples that are not identified within either the primary samples or metastatic samples alone (Figure 3c,d (iv)). Collectively, these MSAI and parallel evolution results suggest that the metastatic samples demonstrate less inter-sample heterogeneity than the primary samples in CRUK0063 and the presence of MSAI and parallel evolution between the primary samples and metastatic samples suggests continued copy number evolution. One area of the genome subject to parallel gain in CRUK0063 is a region of chromosome arm 2p (i), encompassing 2p16, that overlaps the second most commonly amplified locus in lung squamous cell carcinoma revealed by TCGA through GISTIC2 analysis [39,40]. This locus contains the transcription factor *BCL11A,* a known oncogene in triple-negative breast cancer [41] and B-cell lymphoma [42], that has been described as integral to the pathology of lung squamous carcinoma through its interaction with *SOX2* to control the expression of epigenetic regulators [43].

In summary, Refphase revealed novel insights into the evolution of this tumour. The heterogeneity of the primary samples and continued and parallel evolutionary events would have remained hidden without reference phasing and the results from Refphase.

### Refphase reveals previously undetected events in low purity samples

To quantify the extent to which Refphase reveals previously undetected SCNAs, we applied Refphase to multi-sample cohorts from two highly-cited studies: (1) a multi-sample investigation of primary colorectal adenocarcinoma and adenoma [44] (15 tumours, 140 samples), and (2) a matched primary sample and metastatic sample cohort of various primary cancer types and their brain metastases [45] (84 tumours, 196 samples) (Figure 4). These datasets demonstrate a range of copy number landscapes, purity levels, and data types (SNP array and WES) found in both research and clinical studies.

**Figure 4:**
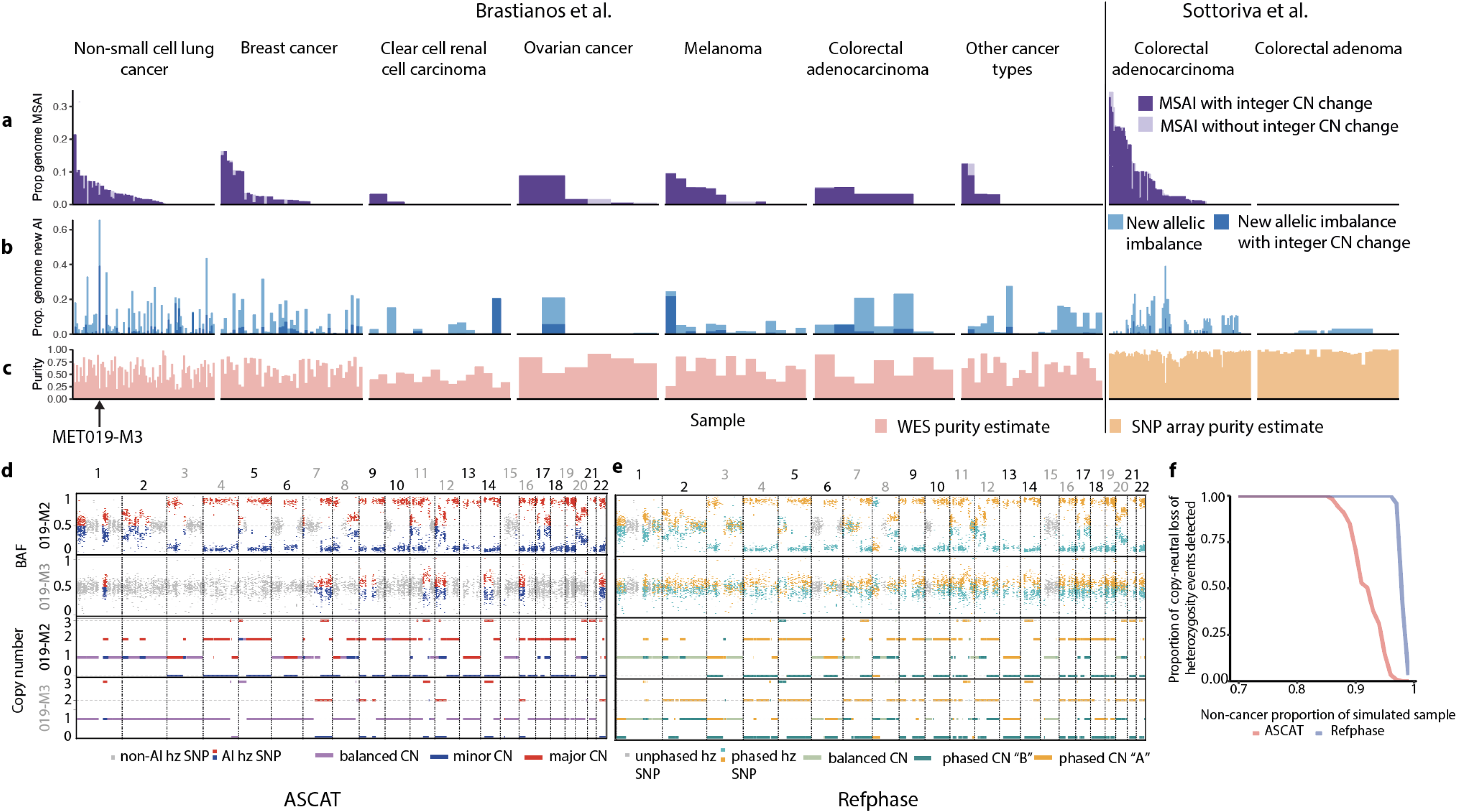
Low cancer cell fraction SCNA detection and cohort-level analysis. **a)** Barplots showing the proportion of the genome affected by MSAI in each tumour sample in the pan-cancer cohort grouped by tumour type. **b**) Barplots showing the proportion of the genome with allelic imbalance that was identified using multi-sample reference phasing and that was previously undetected using ASCAT. Each tumour sample in the pan-cancer cohort is arranged by tumour type. Light blue bars represent the proportion of the genome in each sample affected by previously undetected allelic imbalance that did not result in an alteration in the previously estimated integer allele-specific copy number. Dark blue bars represent instances in which newly detected allelic imbalance that resulted in new integer copy number state being estimated. **c)** Barplots representing the estimated cancer cell fraction of each sample in the pan-cancer cohort grouped by cancer type. **d)** Unphased BAF and integer copy number states across the genome from two samples analysed using ASCAT. **e)** Phased BAF and haplotype-specific copy number states across the genome from two samples analysed using Refphase. **f)** Line plot showing the proportion of simulated copy-neutral loss of heterozygosity events identified at differing non-cancer proportions of a sample using ASCAT (red) or Refphase (blue).

As already described, Refphase’s multi-sample phasing permits the detection of MSAI not otherwise discernible with single-sample data or unphased copy number profiles. Examining our cohort utilising the multi-sample reference phasing of all samples of each tumour, we detected MSAI in 65% of tumours (64/99), affecting up to 34% of the genome (IQR across all tumours [0%,3%]; Figure 4a, Methods). In addition to revealing MSAI, multi-sample phasing can be used to identify previously undetected AI. This newly identified AI is often not MSAI but simply AI with the same major haplotype as the other samples from the same tumour previously identified as demonstrating AI. Refphase detected new AI not previously detected by ASCAT in 77% of tumour samples (256/334). The greatest degree of new AI identified was in sample MET019 in which 66% of the genome was identified to be allelically imbalanced by Refphase which had not been identified as such by ASCAT (Figure 4b).

We observed a significant negative association between sample purity and newly identified AI (Figure 4c; LME coefficient = −0.28 (2.s.f), LME ANOVA p<0.0001, adjusted for patient and cohort - the latter defined by tumour type and profiling platform - as random effects, Supplementary Figure 3a). This result is consistent with Refphase’s ability to rescue low purity samples and is in accordance with previous work demonstrating that SCNA detection using methods designed for single samples is drastically impaired by low tumour purities when sequencing coverage remains unchanged [46]. No significant association was observed between purity and MSAI detection (LME ANOVA p=0.5, adjusted, Supplementary Figure 3b). Schematic examples of the effect of tumour purity and copy numbers states on BAF and LogR profiles are shown in Supplementary Figure 4.

We further explored the relationship between purity and SCNA event detection for the specific case of tumour MET019 from Brastianos *et al.*, a lung adenocarcinoma containing the sample (M3) which demonstrated the highest amount of newly identified AI in our cohort when analysed with Refphase (66% of the genome affected). Notably, this M3 sample showed a markedly lower level of inferred tumour purity compared to the tumour’s other samples (M3 - 21% compared to R1 - 47%, M1 - 80%, M2 - 86%). Using the same phasing that revealed the newly identified AI, we re-estimated the copy number states for all samples across the genome and compared Refphase results to the previous ASCAT-derived estimates. We focussed analysis on detection of CNLOH. ASCAT identified CNLOH affecting five chromosomes in M3 compared to Refphase finding CNLOH on 15 chromosomes, with an increase of 24% of the genome affected when quantified using Refphase. The same analysis of the purer M2 sample yielded an increase of just 0.09% of the genome affected by CNLOH using Refphase (Figure 4d-e). This result supports the previous observation of a negative association between sample purity and the extent of newly identified AI using Refphase.

Finally, to systematically compare ASCAT and Refphase’s ability to resolve AI as a function of purity, we simulated BAF values for CNLOH events at varying tumour purities in samples from multi-sample NSCLC tumours [8] (125 events; each simulated at tumour purities from 1% CCF to 30% CCF at 1% CCF intervals; 200x sequencing coverage) (Methods, Figure 4f). CNLOH events were investigated to ensure that there were no changes to overall sample ploidy and that LogR changes did not influence event detection. Using these simulation parameters, ASCAT was able to detect AI at all simulated CNLOH events at tumour purities of 15% or greater and Refphase at purities of 4% or greater (Figure 4f).

Taken together, multi-sample reference phasing improves the limits of detecting allelic imbalance and offers potentially exciting avenues for improving the sensitivity of SCNA detection at low cancer cell fractions, especially in non-WGS contexts.

### Refphase improves SCNA intra-tumour heterogeneity estimates

Using the same pan-cancer cohort (Figure 5a), we next leveraged Refphase’s relative-to-ploidy SCNA event classification and standardised SCNA intra-tumour heterogeneity quantification (Methods) to explore cancer evolution in a range of cancer types.

**Figure 5.**
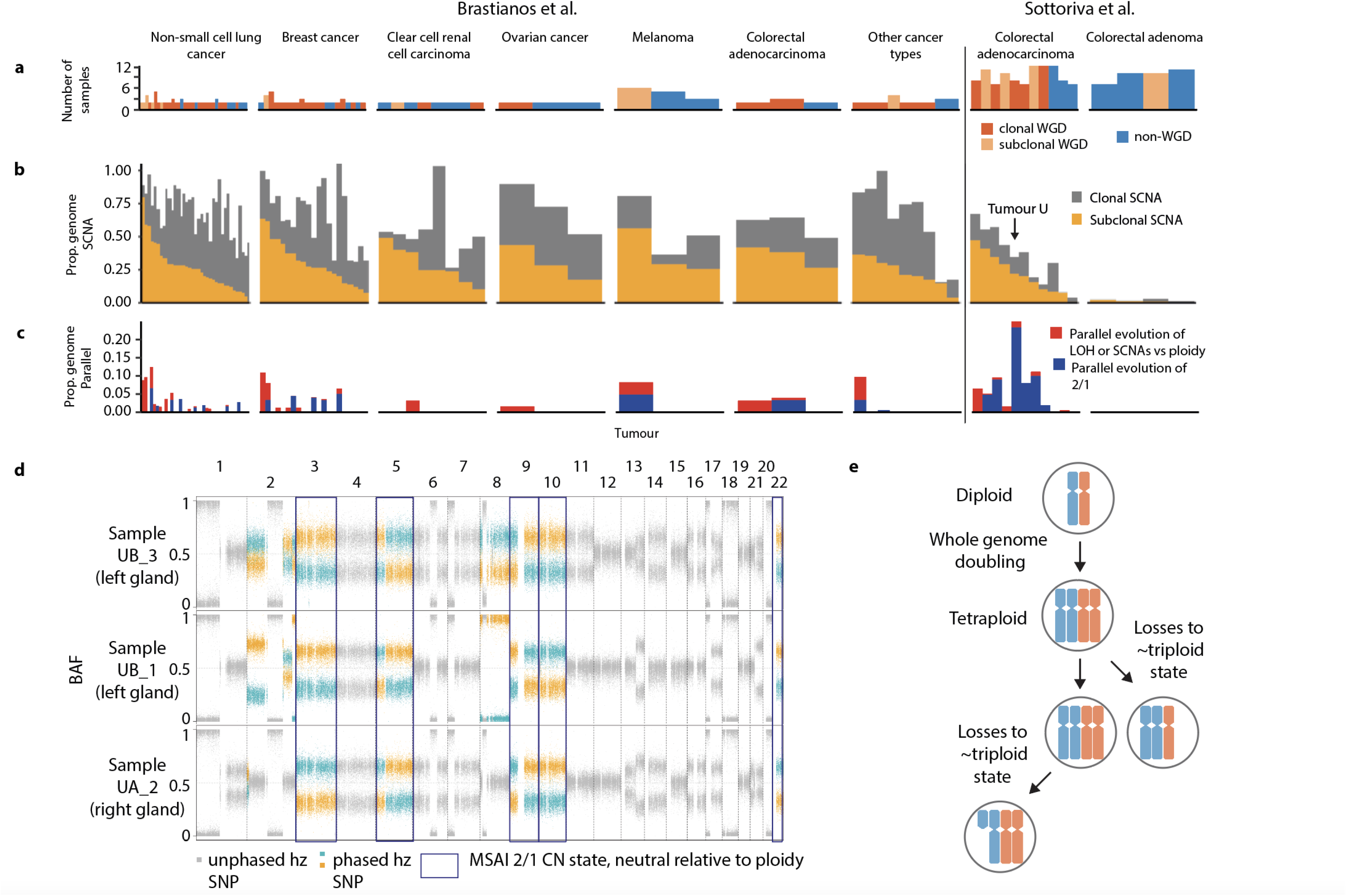
**a)** Barplot showing the number of samples per tumour and colored by WGD clonality status with clonal WGD (dark orange), subclonal WGD (light orange), and non-WGD (blue). **b)** Barplot showing proportion of the genome classified as affected by clonal SCNA (grey) and subclonal SCNA (yellow) from the pan-cancer cohort. **c)** Barplot showing proportion of the genome affected by parallel evolution of SCNAs relative to ploidy or LOH (red) and parallel evolution of 2|1 copy number states (dark blue). **d)** BAF of heterozygous SNPs across the genome from tumour samples from colorectal adenocarcinoma U. Heterozygous SNPs in regions of the genome affected by MSAI are coloured orange and blue according to the phased haplotype they are assigned to. Heterozygous SNPs in regions of the genome unaffected by MSAI are colored grey. Regions of the genome demonstrating parallel evolution of a 2|1 and 1|2 copy number states are highlighted with a dark blue outline. **e)** Schematic demonstrating whole genome doubling and independent subsequent copy number loss events revealed by MSAI.

We first quantified the total proportion of the genome affected by SCNAs (here encompassing relative-to-ploidy gains and losses, and LOH) and the proportion of clonal, early SCNAs, compared with subclonal, late SCNAs (Methods, Figure 5b). We identified clonal SCNAs in every tumour and found that 96% (95/99) of the tumours examined had clonal and 95% (94/99) harboured subclonal SCNAs affecting at least 1% of the genome. A median of 31% of the genome was subject to clonal SCNAs and 25% to subclonal SCNAs meaning that in over half of tumours (52/99), 25% or more of the genome was subject to subclonal SCNAs. The SCNA heterogeneity observed in these 99 tumours of various cancer types supports recent work suggesting that ongoing chromosomal instability is pervasive in cancer [9,10].

Importantly, detection of MSAI at a genomic segment renders any clonal SCNA at that segment subclonal since different haplotypes represent the major allele in different tumour samples. This means that Refphase can resolve additional SCNA heterogeneity which would not be possible with methods relying on unphased data. Notably however, despite the detection of MSAI in a subset of tumours (Figure 4a), the cohort-level effect of multi-sample reference phasing was to decrease the average level of SCNA heterogeneity compared to estimates from ASCAT runs on independently analysed tumour samples (paired Wilcoxon signed rank tests, p<0.05, Supplementary Figure 5). This suggests that segments previously defined as harbouring subclonal SCNAs are now identified to be clonally affected across all tumour samples. This may in part be due to the increased SCNA detection in low tumour purity samples explored in the previous section.

Overall, these analyses highlight how Refphase can provide potentially more accurate overall estimation of SCNA heterogeneity than approaches relying on unphased data.

### Refphase offers new insights into associations between parallel evolution and WGD in a pan-cancer cohort

Whole genome doubling (WGD) is considered a transformative event in tumour evolution [47], and tumours with WGD events show increased prevalence of MSAI [8,9]. In order to explore this relationship in our pan-cancer cohort, we determined WGD status using MEDICC2 run on Refphase output to classify the tumours in our pan-cancer cohort as clonal WGD, subclonal WGD, or non-WGD (Methods, Figure 5a). In keeping with previous results, a higher proportion of the genome was observed to be affected by MSAI in whole genome doubled tumours (clonal and subclonal) than non-WGD tumours (Kruskal-Wallis p=2e-04, Supplementary Figure 6).

We next probed further into the nature of MSAI in our cohort. Specifically, we quantified the extent of parallel evolution of the same type of SCNA event (e.g. gains, losses, LOH) occurring in different samples within the same tumour but affecting different haplotypes, which represents just a subset of MSAI events. Using relative-to-ploidy and LOH definitions of parallel evolution (Methods), we observed parallel evolution in all tumour types in our cohort with the exception of the benign colorectal adenomas (Figure 5c). However, this tumour type also had the lowest levels of SCNAs overall with on average <1% of the genome affected by either clonal or subclonal SCNAs (Figure 5b; median percentage of genome affected: clonal 0.7%, subclonal 0.7%). Notably however, we observed parallel evolution of gains in the Sottoriva *et al.* (malignant) colorectal adenocarcinoma dataset, indicating independent evolution of similar SCNAs in spatially separated areas of tumours, adding additional resolution to the original study [44] (Supplementary Figure 7).

Intriguingly, we observed some tumours, such as colorectal adenocarcinoma U from Sottoriva *et al.* [44] (Figure 5d), with greater than 30% of their genomes affected by MSAI but with only a very small subset of this MSAI constituting parallel gains and losses relative to ploidy (2% of the genome, Figure 5c). Upon exploring MSAI in genomic regions matching the overall ploidy of their respective tumour samples (Methods), we observed that MSAI commonly occurred in a copy-neutral context of arm-level and chromosomal triploidy. Tumour U harboured nine chromosome arms affected by this phenomenon and was found to have undergone a clonal and therefore likely relatively early WGD event. This may suggest parallel evolution of losses from a tetraploid to a sub-tetraploid state (Figure 5e). Consistent with this, 2|1 and 1|2 copy number states were simultaneously observed in 22 tumours, of which 21 were determined to have undergone clonal or subclonal WGD using MEDICC2 (Figure 5a, Figure 5c, Methods) [34]. Other groups have observed such parallel losses in *in vitro* models of WGD [48] or inferred their presence from single-sample data [49], but to our knowledge this is the first time that such independent losses have been observed using phasing in multi-sample data. The parallel evolution of losses from a tetraploid state offers new insights into the potential selective pressure for triploidy and the ability to further determine the timing of such events in tumour evolution.

Together, these results highlight the importance of leveraging information from multiple samples to quantify SCNA heterogeneity during tumour evolution and demonstrate the technical advances offered by Refphase compared to single-sample copy number callers, including its ability to identify MSAI, its increased sensitivity of CNLOH detection, and its potentially more accurate overall estimation of SCNA heterogeneity.

## Conclusions

We have shown that Refphase provides phasing of heterozygous SNPs into long-range haplotypes, allows the identification of SCNA-mediated parallel evolution and MSAI [8], and improves the limits of detection of AI in low purity samples.

The long-range haplotypes derived by Refphase augment existing haplotype-specific approaches for copy number calling in single-cell DNA [29] and RNA [30,50] sequencing technologies, and are distinct from haploblocks produced by population-level statistical phasing approaches [25,51–53] or those derived from chromatin structure data [54,55] or long read sequencing [56,57]. Reference-phasing haploblocks are not limited in length by recombination rates, read lengths, or structural constraints, and instead stretch the full length of the evolutionary gain or loss event that gave rise to the AI, frequently a whole chromosome or chromosome arm. Refphase haploblocks have the advantage that all variant alleles within regions of AI with sufficient sequencing depth can be assigned to their haplotype-of-origin. Refphase is therefore able to phase rare and private variants or those from understudied ethnic groups for whom reference sets of haplotypes are unavailable, doing so at a fraction of the computational cost of statistical phasing approaches, and with broad applicability to a variety of different experimental techniques, including WGS, WES, SNP arrays and targeted sequencing approaches, and without reliance on external databases.

Besides haplotype reconstruction and phasing of SCNAs, Refphase offers a standardised characterisation and quantification of SCNA intra-tumour heterogeneity from bulk multi-sample tumours, where the field previously relied on simple metrics such as the weighted genome instability index [58] or fraction of genome altered [59]. In the context of multi-sample sequencing, only a few algorithms utilise data from multiple samples for SCNA estimation and either do not produce haplotype-specific copy number estimates [60,61] or use less powerful statistical phasing limited to WGS [62] in concert with reference haplotype databases. Refphase is broadly applicable, supporting multiple formats of user-provided single-sample copy number segmentations as input, including those from commonly used copy number callers such as ASCAT [22], to provide reproducible estimates of SCNA intra-tumour heterogeneity that allow comparisons both between datasets and within grouped sets of samples within a patient’s disease, for example to contrast primary samples with metastases. Downstream integration with MEDICC2 [34] also permits standardised detection of WGD.

It is through this joint analysis of samples from a single tumour, utilising WGD detection and Refphase’s relative-to-ploidy classifications, that we observe previously undetected copy-neutral MSAI of arm-level and chromosomal triploidies in WGD tumours. This finding, indicating parallel evolution from a tetraploid to a sub-tetraploid state, may suggest selective pressure for triploidy and offers a new avenue for the exploration of tumour copy number evolution. Additionally, the detection of MSAI and parallel events using reference phasing indicates a hitherto underappreciated number of independently occurring SCNAs.

Despite its many advantages, our method is not without its limitations. Refphase chooses a single best reference sample to inform phasing for any given bin meaning that the same heterozygous SNPs are assayed across multiple samples from the same tumour; however, each sample with AI may provide useful phasing information. While Refphase characterises inter-sample heterogeneity by determining the presence and absence of SCNAs in each tumour sample, it does not attempt to identify within-sample subclonal SCNAs present in only a subset of cells in a single sample, in contrast to tools such as TITAN [63] and Battenberg [27] designed for use with WGS. Refphase also does not attempt to identify subclonal clusters of co-occurring SCNAs present across multiple samples [61]. Additionally, while Refphase updates ploidy estimates of each sample as it performs its phasing, it is reliant on robust initial estimates of purity, ploidy, and input segmentation.

The SCNA heterogeneity and parallel evolution revealed and characterised by Refphase is indicative of ongoing chromosomal instability in the tumours examined. However, it should be noted that this is likely still an underestimate of the actual ongoing chromosomal instability present in these tumours, as only a small proportion of each tumour is sequenced [64]. Whilst no significant correlation was observed between the number of samples per tumour and SCNA heterogeneity in this pan-cancer cohort (LME ANOVA p=0.1, adjusted, Supplementary Figure 8a), further analysis in cohorts with a greater range of per-tumour sample numbers and tumour types is required. Additionally, whilst Refphase’s ability to infer SCNA events at low tumour purities exceeds that of single-sample copy number calling methods (Figure 4f), even with Refphase, we observe a moderately significant association between the range of purities within a tumour (‘Tumour Purity Difference’) and the degree of SCNA subclonality (LME coefficient = 0.28, LME ANOVA p=0.04, adjusted, Supplementary Figure 8b), indicating that tumour purity may interfere with the estimation of SCNA clonality. Finally, while multi-sample reference phasing reveals instances of parallel evolution from distinct haplotypes, parallel evolution from the same haplotype cannot be detected and as such the amount of parallel evolution found in this cohort may be an underestimate. Detection of such parallel events from the same haplotype instead can be resolved with phylogenetic methods which allow for multiple mutations of the same site, such as MEDICC2 [34].

Despite these limitations, no other tool provides phasing, detection of AI in low purity samples, estimation of copy number states and parallel evolution, and systematic characterisation of SCNA heterogeneity.

In the future, combining statistical population-based phasing with multi-sample reference phasing will further strengthen this approach, in particular for genomic regions with weak AI. As multi-sample bulk DNA sequencing data of tumours and their metastases become increasingly common, new opportunities for improving our understanding of tumour evolution and how it relates to prognosis and response to treatment, will arise. Algorithms such as Refphase that are able to leverage such data to quantify mutations and their intra-tumour heterogeneity across a patient’s disease will be vital to support new insights to inform care for cancer patients.

## Methods

### Nomenclature

We report integer allele-specific copy numbers from SCNA estimation tools that do not perform either statistical or multi-sample reference phasing in the major/minor configuration. We use the / symbol to separate the most common allele and least common allele for a copy number segment or bin covering a genomic region. For example, in two samples (S1 and S2) from the same tumour in which we observe the same unphased allele-specific integer copy number state of 2 of one allele and 1 of the other allele at the same genomic region, we report the allele-specific integer major and minor allele copy number estimate of this bin as 2/1 in S1 and 2/1 in S2. We would also report the non-allele specific total copy number in sample S1 as 3 and in sample S2 as 3.

In the context of multi-sample reference phasing as performed by Refphase, we report phased haplotype-specific “A” haplotype and “B” haplotype copy number written in the format haplotype “A” **|** haplotype “B”. If Refphase identified MSAI between our example S1 and S2, maintaining the total copy number states of 3 in both, with haplotype “B” being present at 1 copy in S1 but at 2 copies in S2, we would report the copy number in S1 as 2|1 and in S2 as 1|2.

### Refphase input data requirements

To perform multi-sample reference phasing for a tumour with ***N*** samples, an initial single-sample copy number segmentation and initial estimates of tumour purity ρ_*j*_ and ploidy ψ_*j*_ must be obtained as input for Refphase by applying a single-sample SCNA calling algorithm to each tumour sample *j* ∈ {1 .. *N*} independently.

Single-sample tools typically follow a common approach for the detection of allele-specific SCNAs, as defined in [22]. Briefly, for each tumour sample, sequencing reads are aggregated at heterozygous germline variants in each tumour and paired normal sample over both parental alleles, yielding two readouts: the log ratio *L*_*i*_ of read counts at variant *i* compared to the matched normal (LogR) and the relative frequency of the minor (B) allele read counts over the total read counts (B-allele frequency, BAF) *B*_*i*_ at variant position *i*. SCNAs are called by segmenting both BAF and LogR tracks into homogeneous segments and determining fractional or integer copy numbers for each segment, jointly inferring the purity ρ_*j*_ of sample *j* (the fraction of cancer cells over total number of cells) and the average ploidy ψ_*j*_ of the tumour sample *j* as free parameters [22]. Copy numbers are determined per allele, but due to unknown phasing of the underlying germline variants, the values are reported as major (larger) and minor (smaller) copy number instead.

Refphase currently directly supports input derived from the popular segmentation algorithms ASCAT [22] and Sequenza [31], but will run on any user-supplied initial segmentation result which includes estimates of: allele-specific copy number segments (genomic positions and major and minor allele copy number states); sample purity and ploidy estimates; log ratios (LogR) and B-allele frequencies (BAFs) of single nucleotide polymorphisms (SNPs); and - for non-ASCAT input - SNP heterozygosity annotations, for each of the ***N*** tumour samples (***N***≥2) and a matched normal from the same patient.

### Refphase algorithm overview

Refphase achieves long-range phasing and haplotype-specific estimation of SCNAs through application of the multi-sample reference phasing algorithm. We assume that the input purity estimates for each tumour sample are correct and Refphase utilises these, alongside the LogR and BAF, to characterise SCNA heterogeneity of the genome in ***m*** bins of variable sizes. These bins are derived from a minimum consistent segmentation (see below) created from the input copy number segments for each tumour sample.

The four outputs from Refphase are: (1) a phasing of heterozygous SNPs, (2) an updated set of phased fractional and integer copy number states across the genome for each sample, (3) a sample-level summary of SCNA events, and (4) a summary of SCNA clonality and intra-tumour heterogeneity, defined either at tumour level or between and across user-defined subgroups of samples.

For a schematic overview of the Refphase algorithm see Figure 1.

### Minimum consistent segmentation

Having internally preprocessed input data to a standard Refphase format, Refphase combines individual single-sample segmentations to generate a minimum consistent segmentation (MCS) for the set of all ***N*** samples of a tumour (Figure 1a). To do this, Refphase first defines the combined set of breakpoints as the union of the set of individual breakpoints, keeping track of the samples-of-origin. Of the new set of segments defined by this union of breakpoints, only segments present in all samples are retained in the subsequent analysis. An iterative merging strategy is then employed which merges breakpoints that originated from different samples if their pairwise distances are below a user-defined maximum gap threshold (default = 100kbp). These slight variations in breakpoint position typically result from variability in the estimation of breakpoint positions in individual samples even if the true underlying breakpoint is the same. The restriction to only merge breakpoints that originated from different samples of origin preserves focal amplifications and losses present prior to merging in individual samples and yields a final set of **m** bins.

### Reference sample selection

Once the MCS is established, Refphase employs the multi-sample reference phasing algorithm to achieve long-range phasing of germline variants and assignment of SCNAs to haplotypes (Figures 1b-e). Multi-sample reference phasing leverages the fact that the phase of the underlying germline genetic variants is constant between samples from the same patient. In a segment affected by allelic imbalance, variants whose alternative alleles are residing on the chromosome with higher copy number will show a theoretical BAF above the segment mean, whereas those residing on the minor copy number chromosome will show BAF values below the segment mean, effectively providing phasing information about the variants contained in the segment.

To leverage this information, for each MCS segment, Refphase first iterates through all segments and samples and, for those segments determined to have a bimodal BAF profile in at least one sample, Refphase assigns the sample with the highest mirrored mean BAF as the reference sample (Figure 1b), where the mirrored mean BAF is defined as:

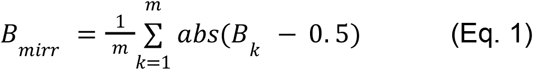

where *B_k_* is the BAF at variant position *k*; *m* is the total number of variants within the segment indexed from 1; and where *abs()* returns the absolute value (≥0) of the expression contained in brackets.

Segments without AI in at least one sample in the input are not considered for reference phasing.

### Haplotype phasing of reference sample segments

Within each reference sample, Refphase then assigns alternative alleles to haplotypes *H_i_* by comparing the empirical BAF of each variant *i* against the segment mean (Figure 1c):

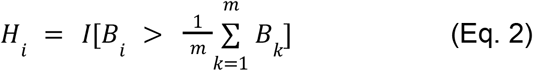

where *B_i_* is the BAF at variant position *i*; *m* is the total number of variants within the segment indexed from 1; and *I* is an operator assigning each variant position to one of two haplotypes based on the evaluation of the logical expression in square brackets.

### Reference phasing across non-reference sample segments

In the next step, Refphase applies this phasing information to the variants in the same segment in all other non-reference samples to determine haplotype-specific BAF values for every other sample (Figure 1d).

### Haplotype-specific copy number quantification

After the variants have been assigned to haplotypes, Refphase uses haplotype-level BAF and LogR values for re-estimation of haplotype-specific copy numbers (Figure 1e).

Here, each sample is tested for AI using a Wilcoxon rank-sum test between the BAF values of each haplotype (5% family-wise error rate) and the effect size of AI is determined using *Cohen’s d* [65]. If additional AI is detected compared to the initial copy number states, or if the option is applied universally by the user, copy numbers are re-estimated using either a default parametric or a non-parametric model for each haplotype separately. The parametric model estimates the new haplotype-specific copy number *n_i_* in segment *i* based on the mean BAF *B_i_* and LogR *L_i_* of that segment as well as sample purity ρ and average tumour ploidy ψ obtained from the initial segmentation as follows [22]:

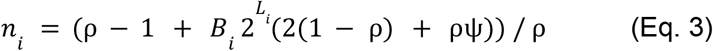

The alternative non-parametric model employs a Gaussian naive Bayes classifier to predict a haplotype-specific copy number configuration from the mean BAF and LogR values of a segment. To train the model, Refphase uses the mean BAF and LogR values of all other segments of the same sample and from the same segment in all other samples as training data to account for the typical BAF/LogR distributions of the current sample as well as those values from homologous segments in other samples. After re-estimation of all copy number segments, the average tumour ploidy ψ of each sample is re-calculated as the sum over the total copy numbers of all segments *i* = 1.. *m* weighted by their genomic width *w_i_* (in bp) relative to the total width of the genome (in bp).

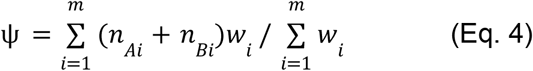

Re-estimated fractional and integer copy numbers are available to the user.

### Horizontal phasing

After individual segments have been phased across all samples, there is still no phasing relationship between neighbouring segments ‘horizontally’ along the genome. We employ a method for phasing along the genome that uses a parsimony assumption, i.e. the data can be explained with the least amount of copy number changes between neighbouring segments (Figure 1e). For this we compare every segment with its preceding segment and flip the haplotype assignment across all samples such that the overall Hamming distance between neighbouring segments is minimal (Supplementary Figure 1). The overall Hamming distance of segment *i* is defined as

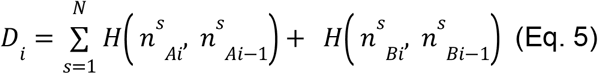

where we sum over all ***N*** samples and calculate the Hamming distance *H* between the copy number and its predecessor for both haplotypes *A* and *B*. The Hamming distance *H*(*x*, *y*) computes to 1 if *x* ≠ *y* and 0 otherwise. Flipping the haplotypes of segment *i* amounts to exchanging *n^s^_Ai_* and *n^s^_Bi_* for all samples *s*. If flipping the haplotype assignment of segment *i* does not result in a change of distance *D*_*i*_ (i.e. *D*_*i*_ = *D^flipped^_i_*), the segment *i* is compared to its next predecessor (segment *i-2*). This is repeated until *D_i_* ≠ *D^flipped^_i_* or the beginning of the current chromosome is reached.

As the assignment to haplotype A and B is arbitrary across chromosome boundaries, for plotting purposes, we flip the A and B allele assignment for all samples on a chromosome-by-chromosome level to ensure that on average haplotype A has a higher copy number than haplotype B.

### Sample-level SCNA calling

Following reference phasing, Refphase uses the re-estimated sample ploidies, input purities and segment-level LogR data to call SCNA events in each sample (Figure 1f).

We consider an SCNA to be a deviation of any length from the diploid major/minor copy number state of (1,1). SCNAs with an unequal number of copies on both parental alleles (corresponding to a significant deviation of the BAF from its balanced value of 0.5) are termed allelic imbalances (AI).

Specifically, Refphase calls segment *amplifications*, *gains* and *losses* relative to ploidy as well as *LOH* events and *homozygous deletions*. Events are called for each segment and sample independently of others by comparing the mean segment LogR distributions to calculated purity-ploidy derived event thresholds (Equations 6–8).

The >2× ploidy threshold is the same threshold used for clinical decision making in HER2+ breast cancer using fluorescence in situ hybridization samples [37].

Amplifications are called if

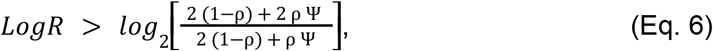

gains are called if

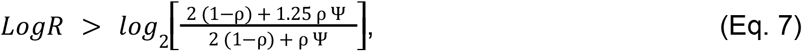

losses are called if

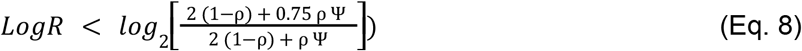

with sample purity ρ and sample ploidy Ψ.

By default, Refphase also provides additional versions of sample-level SCNA event calls aside from comparing mean segment LogR values to the calculated thresholds (Eq.6–8). In another version, LogR values within a segment are compared to the calculated thresholds (Eq.6–8) using a one-tailed Student’s t-test, as in [9]. Additionally, a diploid reference can also be used for both mean and t-test approaches rather than purity-ploidy derived event thresholds. All versions of SCNA calls are available to the user in summary Refphase objects after the completion of the core Refphase algorithm, and plotting options are available to visualise the preferred SCNA event output graphically. For Figures 2 and 3 and associated proportion-of-genome metrics, mean segment LogR values were compared to purity-ploidy derived event thresholds.

LOH events are called for segments in which the rounded copy number state of one allele is 0 and the other strictly greater than 0. Homozygous deletion events are called for those segments where both alleles within a sample have rounded copy number state of 0. Copy neutral LOH (CNLOH) is called for segments in which the major rounded integer copy number state equals the rounded tumour sample ploidy.

### Patient-level SCNA calling

Finally, a tumour-level event summary is calculated (Figure 1g).

Inter-sample heterogeneity is quantified. The presence or absence of each class of relative-to-ploidy event, loss of heterozygosity (LOH), and homozygous deletions (HDs) in each tumour sample for each minimum consistent segment is noted. This presence or absence classification is then examined in the context of MSAI detection to determine whether each event affects the same allele in all samples.

For a given segment, an SCNA is considered to be clonal if it is present in every sample and affects the same allele in all samples. An SCNA is assigned as subclonal if it is present in at least one sample but simultaneously absent in at least one other sample (inter-sample subclonality). Crucially, in cases in which the same relative-to-ploidy or LOH event type is determined to occur in every sample of a given tumour but in the context of MSAI such that a different allele is deemed to be the major allele in different samples at the same segment, the event is assigned as subclonal and parallel and not as clonal, since the SCNAs in different samples are deemed to be of different origin and affecting different haplotypes. Specifically, a mirroring of alleles must be observed between the specific samples harbouring the event type of interest for the event to be called parallel. For example, in a tumour composed of three samples, should a gain SCNA be called in Samples 1 and 2 only, the major alleles must differ in Samples 1 and 2 for the gain event - already subclonal - to be called parallel, regardless of allelic arrangement in Sample 3.

Having assigned individual SCNAs as clonal or subclonal, the proportion of the genome affected by each specific event type (e.g. clonal relative-to-ploidy gains, parallel LOH, etc.) is calculated. Proportion of genome measures are calculated over the sum of minimum consistent segment widths for a tumour.

Plotting options are available to visualise tumour-level summary metrics and the degree of reassignment of events from clonal to subclonal based on MSAI context.

### Refphase user-defined grouping functionality

Refphase also offers the option for a user to define subgroups of samples and calculate group-level (instead of whole-tumour-level) summary metrics and create within- and between-group summary plots. Examples of primary-sample-specific and metastases-sample-specific summary tracks in which clonality and heterogeneity analyses are restricted to the respective subgroups of samples are shown in Figure 3.

### Refphase plotting functionality

Figures 2 and 3 and Supplementary Figure 2 showcase selected tracks from Refphase across-genome plots. Refphase provides several optional output plots as standard including across-genome, chromosome-level, user-defined sample subgroup-oriented, and tumour-level event summary plots.

### MEDICC2 implementation and whole genome doubling detection

The phylogeny showcased in Figure 3 was generated by applying MEDICC2 (version 0.6b1) [34] to re-estimated integer copy number states derived from reference phasing of the named 10 input samples for CRUK0063 using default parameters.

In order to derive the WGD status for the Brastianos *et al.* [45] and Sottoriva *et al. [44]* datasets, MEDICC2 bootstrapping WGD detection was used. In short, we create 100 bootstrapping datasets by resampling the data chromosome-wise (i.e. drawing 22 chromosomes with replacement) and check whether MEDICC2 detects a WGD. If at least 5% of the bootstrap runs detect a WGD, the sample is labelled as WGD-positive. This low threshold offsets the otherwise conservative WGD detection.

### Definitions of newly identified AI and CNLOH (Figure 4)

ASCAT [22] AI was defined at sample-level and assigned for segments where ASCAT non-integer copy number was not equal for the two alleles. Refphase AI was assigned for segments at sample-level using the previously described methods (Methods - Haplotype-specific copy number quantification). CNLOH was assigned to segments in which the rounded major copy number state equalled the rounded sample ploidy and the rounded minor copy number state equalled 0, using copy number states and ploidies from ASCAT and Refphase accordingly. Newly identified AI and CNLOH specifically referred to scenarios in which the respective event had not been identified in ASCAT and was identified using Refphase. For cases in which multiple ASCAT segments overlapped with the Refphase segment being assessed for AI or CNLOH, data for the ASCAT segment most overlapping the Refphase segment under investigation was used.

### Copy-neutral LOH event simulation (Figure 4)

CNLOH events were simulated in chromosomal segments from NSCLC multi-sample bulk sequencing from the TRACERx100 cohort [8].

First, Refphase was run on the TRACER×100 tumour samples. Then, pairs of samples were selected from tumours where at the same genomic region, defined by a Refphase bin, one sample, referred to as “reference”, demonstrated AI allowing a reference phasing to be obtained and the other sample, referred to as “test” demonstrated total copy number equal to the overall sample ploidy with no AI. Specifically, for a genomic bin, the reference sample demonstrated AI if the condition in Equation 9 was satisfied:

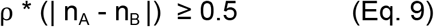

where ρ is the tumour purity, n_A_ the copy number state of allele A and n_B_ the copy number state of allele B.

Additionally, for candidate samples and segments to be chosen, the following conditions must also be satisfied: Genomic bins had to be ≥1/5 of the size of the chromosome on which they were located and contain ≥1/5 of the heterozygous SNPs present on the same chromosome; the ploidy of the test sample had to be between 1.8 and 2.2 or between 3.8 and 4.2 (lower and upper bounds inclusive); and the total rounded copy number in the genomic bin should equal the total rounded ploidy of the test sample.

This candidate segment selection approach produced 125 segments that were then used to simulate CNLOH at the genomic region in the test sample. CNLOH was simulated at various cancer cell fractions (CCFs) of the sequenced sample, ranging from 1% to 30% in steps of 1%. Specifically, BAF values were assigned at each of the heterozygous SNP positions in candidate segments using a binomial distribution (R rbinom function) with simulated probability equal to the mean BAF which would be predicted for each phased allele at the segment based on copy number and simulated purity and with the number of trials equalling a simulated sequencing coverage of 200(x).

These simulated BAF profiles are then used as input to ASCAT allele specific piecewise constant fitting segmentation and Refphase, making use of the phasing derived reference sample, and detection of AI reported for each method independently for each of the 125 candidate segments at each simulated CCF value. Detection of AI was reported as detection of a CNLOH event.

### Initial copy number estimates for data types

#### SNP Array

Tumour cellularity and ploidy for each sample assayed with SNP arrays were estimated using the ASCAT algorithm [22]. ASCAT was then used to identify SCNAs that were provided as the initial input for further clonality analysis. One dataset included in our cohort consists of SNP array data [44]. Processed LogR and BAF values as generated in the original papers were obtained from the GEO database. Copy number analysis was performed using ASCAT v2.3 using default parameters set to 1 for sequencing data [22].

#### Whole exome sequencing

All datasets downloaded were processed from FASTQ files using a previously described pipeline [8,9]. Tumour cellularity and ploidy for each sample assayed with exome sequencing were estimated using ASCAT [22] and these estimates as well as the copy number segmentation were taken forward for analysis with our multi-sample SCNA clonality approach.

## Tables

Supplementary Table S1 - Cohort Overview for Tumours Analysed in Main Figures

## Declarations

### Availability of data and materials

Refphase is available as a user-friendly standalone R package from http://bitbucket.org/schwarzlab/refphase. All code relating to this publication is also available at Zenodo https://doi.org/10.5281/zenodo.7148458. All datasets used in this study can be found in Supplementary Table S1. Clinical trial information (if applicable) is available in the associated publications.

## Authors’ contributions

RFS, PVL, NM, and TBKW conceived the project; RFS, NM, PVL, TBKW, MRH, TLK, and ECC designed the method; MRH, ECC, TLK, TBKW, NM, and RFS implemented the method; TBKW and ECC processed and analysed the TRACERx non-small cell lung cancer copy number data; TBKW, ECC, and ELL processed and analysed the Brastianos and Sottoriva datasets copy-number data; TLK performed whole genome doubling estimation; TBKW implemented the copy number simulations; TBKW, RFS, ECC, TLK, NM, KH, PVL, and CS wrote the manuscript.

## Acknowledgements

TLK and RFS kindly thank BIFOLD and Klaus-Robert Müller for support.

## Supplementary Figures

**Supplementary Figure 1:**
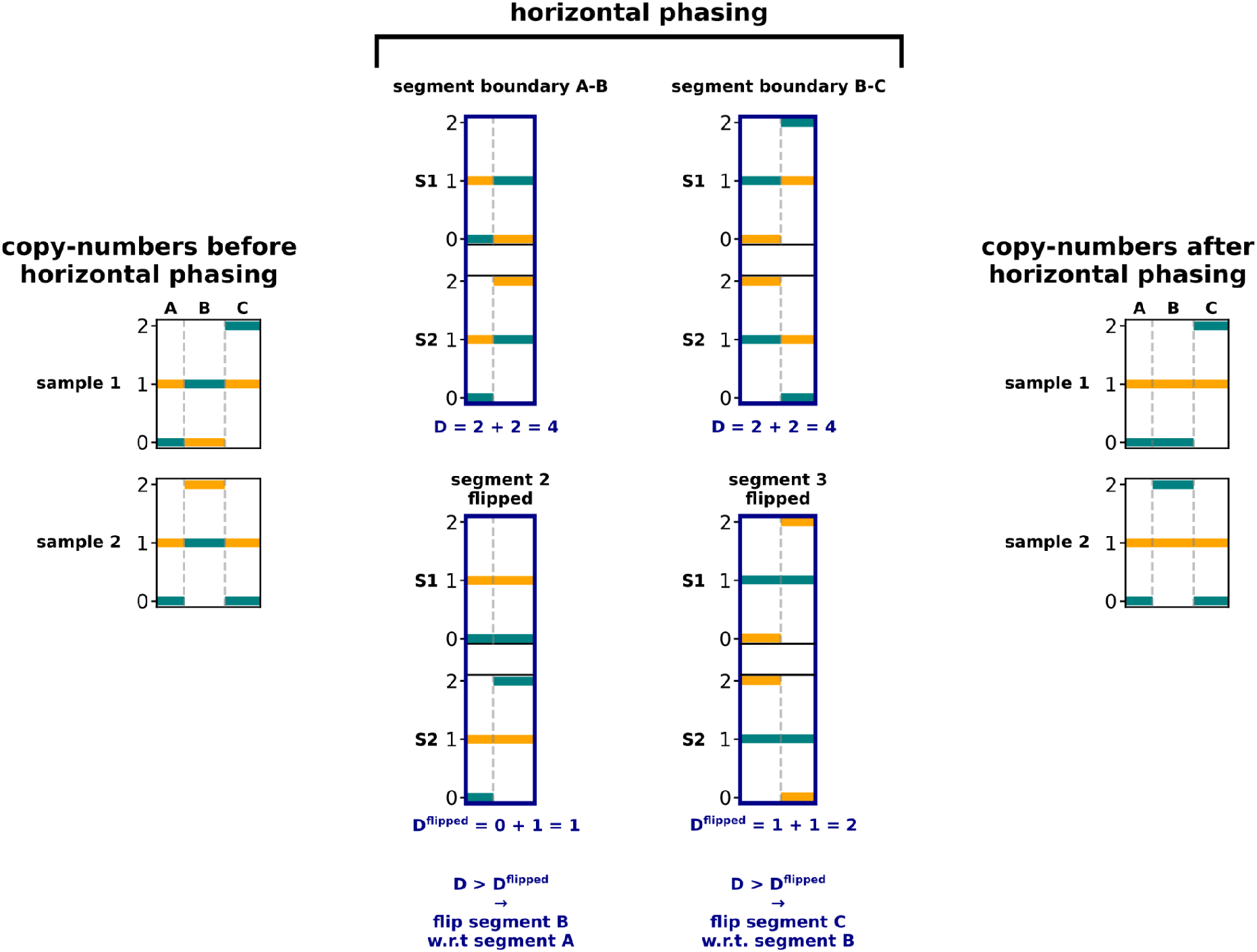
Workflow of the horizontal phasing algorithm. Adjacent segment or bin boundaries are evaluated in both phasing configurations (standard, or “flipped”), where any haplotype “flips” are maintained across all samples from the tumour. Distances between a diploid normal and both configurations are calculated to determine which configuration is optimal. Ultimately this yields a final phased copy number profile with minimal distance to a diploid normal.

**Supplementary Figure 2:**
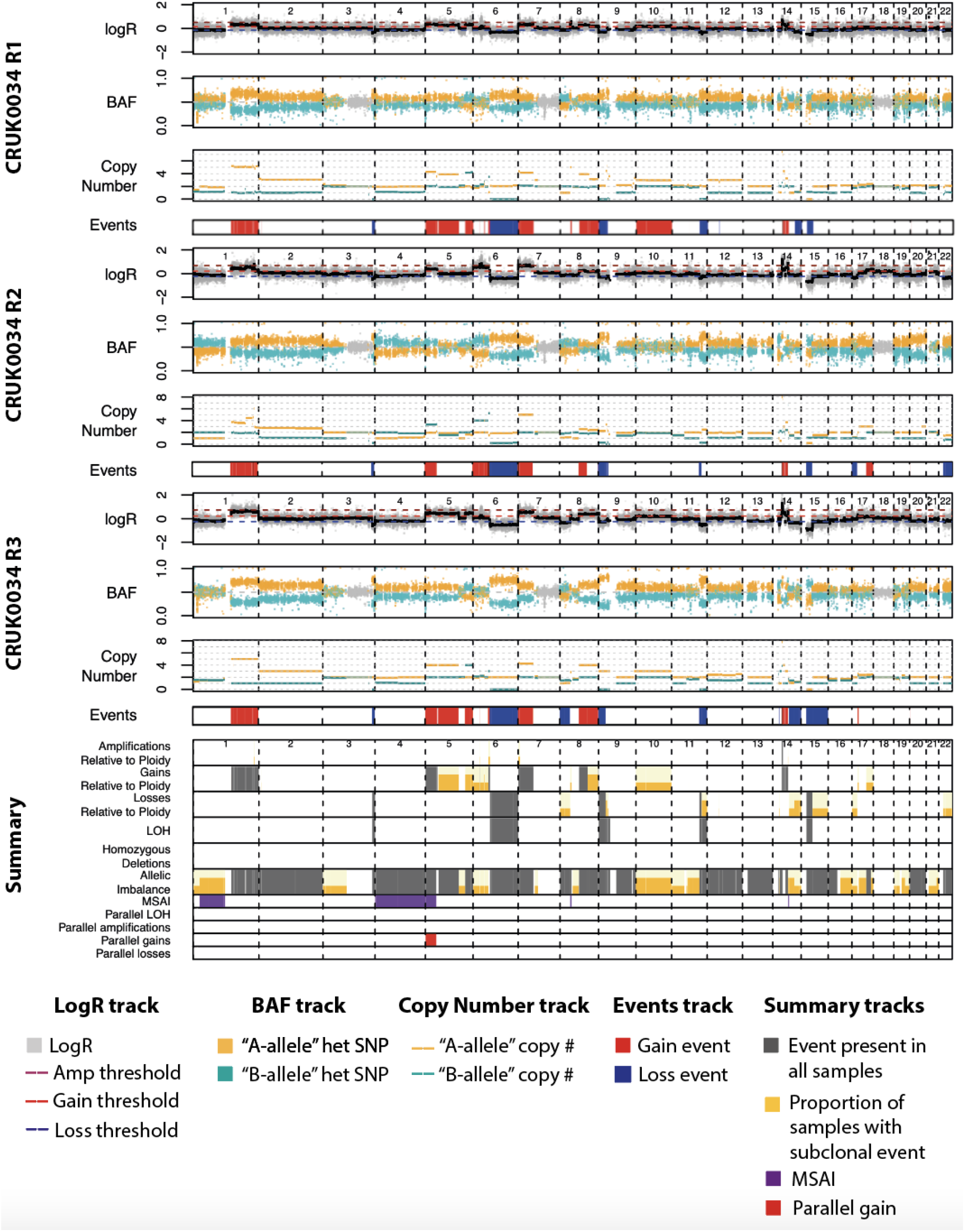
Full Refphase across-genome plotting output for CRUK0034. The upper three panels show tracks for log read-depth ratio (LogR), B-Allele Frequency (BAF), re-estimated fractional copy number states, and somatic copy number aberration (SCNA) event calling at a sample level. The bottom panel (‘Summary’) gives a tumour-level summary of SCNA event clonality and detection of mirrored subclonal imbalance (MSAI), loss of heterozygosity (LOH) and parallel events. Events are called relative-to-ploidy using a mean logR threshold.

**Supplementary Figure 3:**
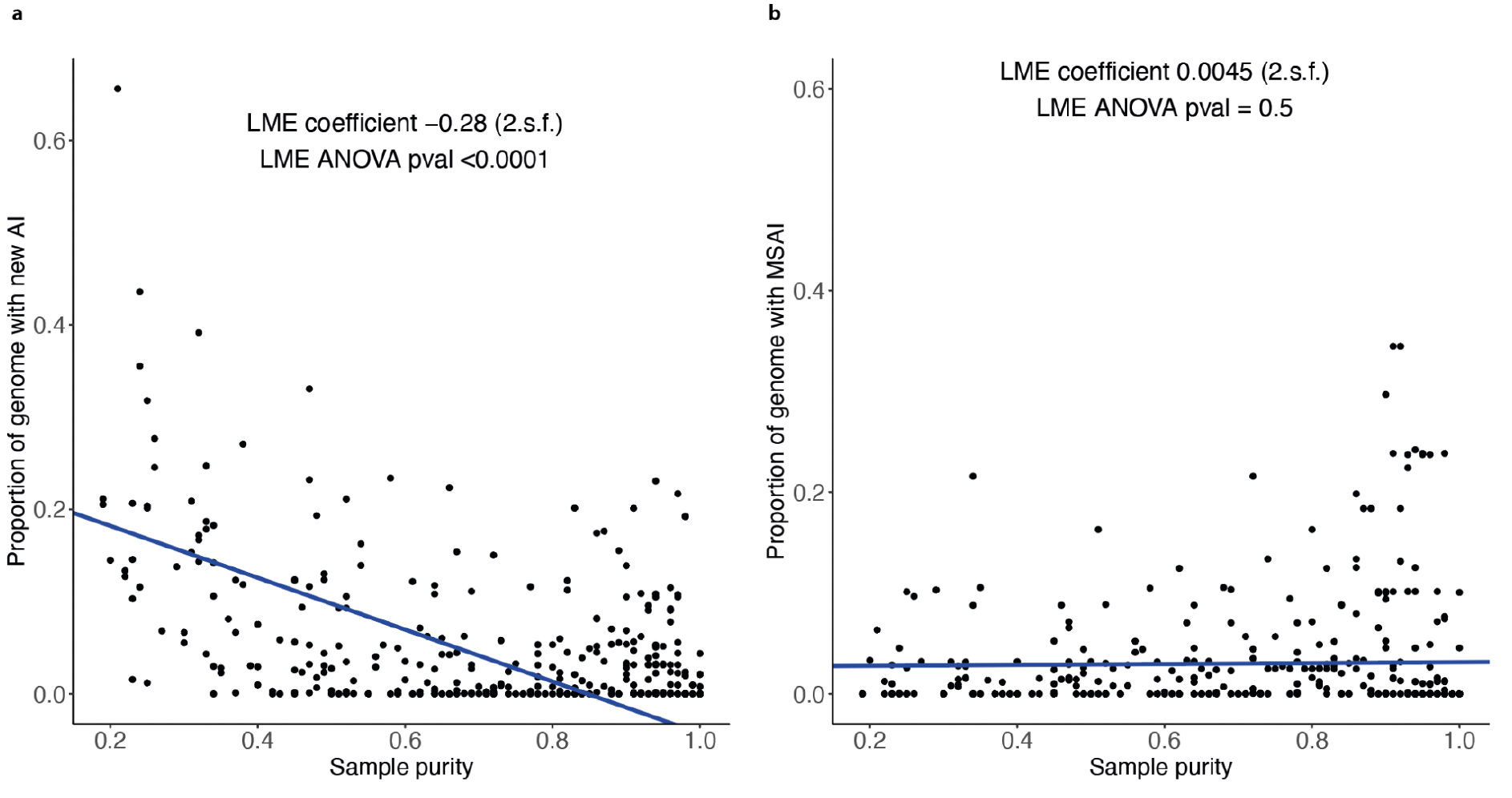
Associations with sample purity. The association between sample purity and **(a)** proportion of genome with newly identified allelic imbalance (LME coefficient = −0.28 (2.s.f.), LME ANOVA p<0.0001), and **(b)** proportion of genome with MSAI (LME coefficient = 0.0045 (2.s.f.), LME ANOVA p=0.5). Proportion of genome data is calculated using Refphase. Analyses are undertaken for the 336 tumour samples from 99 tumours in the pan-cancer cohort described in Figure 4 and summarised in Supplementary Table S1. Linear mixed effect (LME) coefficients and ANOVA p-values shown are adjusted for patient and study cohort (defined by tumour type and profiling platform) as random effects, calculated using the nlme R package and maximum likelihood method. Best fit lines shown are derived using the LME model coefficient and intercept values.

**Supplementary Figure 4:**
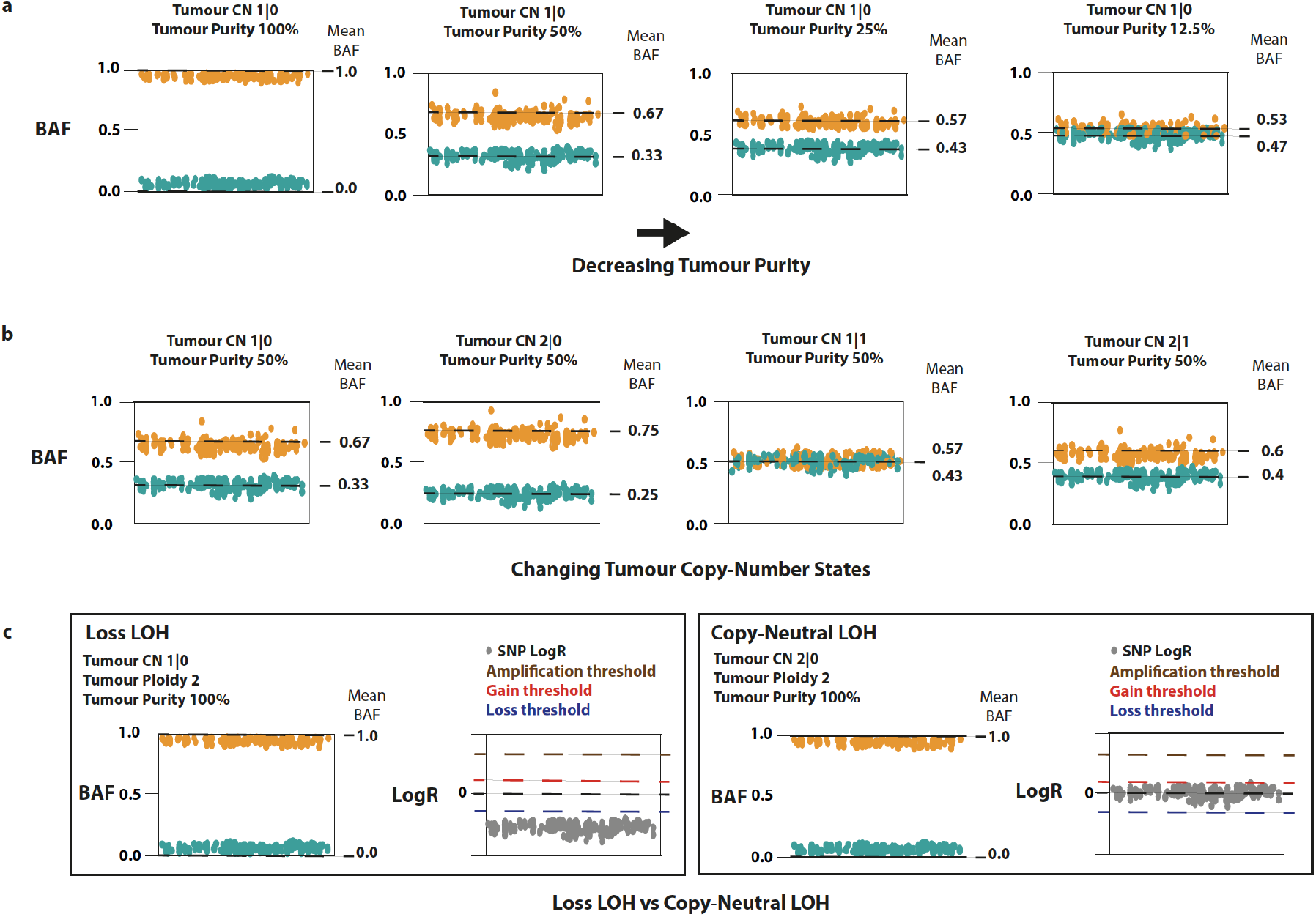
The effects of tumour purity and copy number states on B-Allele Frequency (BAF) and log read-depth ratio (LogR) profiles. **a)** An example of the effect of decreasing tumour purity on BAF band separation. **b)** The effect of varying copy number states on BAF band separation for a tumour of fixed purity, here 50%. **c)** Example of typical BAF and LogR profiles for a loss of heterozygosity (LOH) with total copy number loss (left) and copy-neutral LOH (CNLOH) (right). Relative-to-ploidy thresholds are indicated by dashed lines and are derived for a diploid 100% pure tumour using the formulae described in the Methods of this manuscript. Orange and blue points throughout plots represent the phased “A” haplotype and “B” haplotype respectively.

**Supplementary Figure 5:**
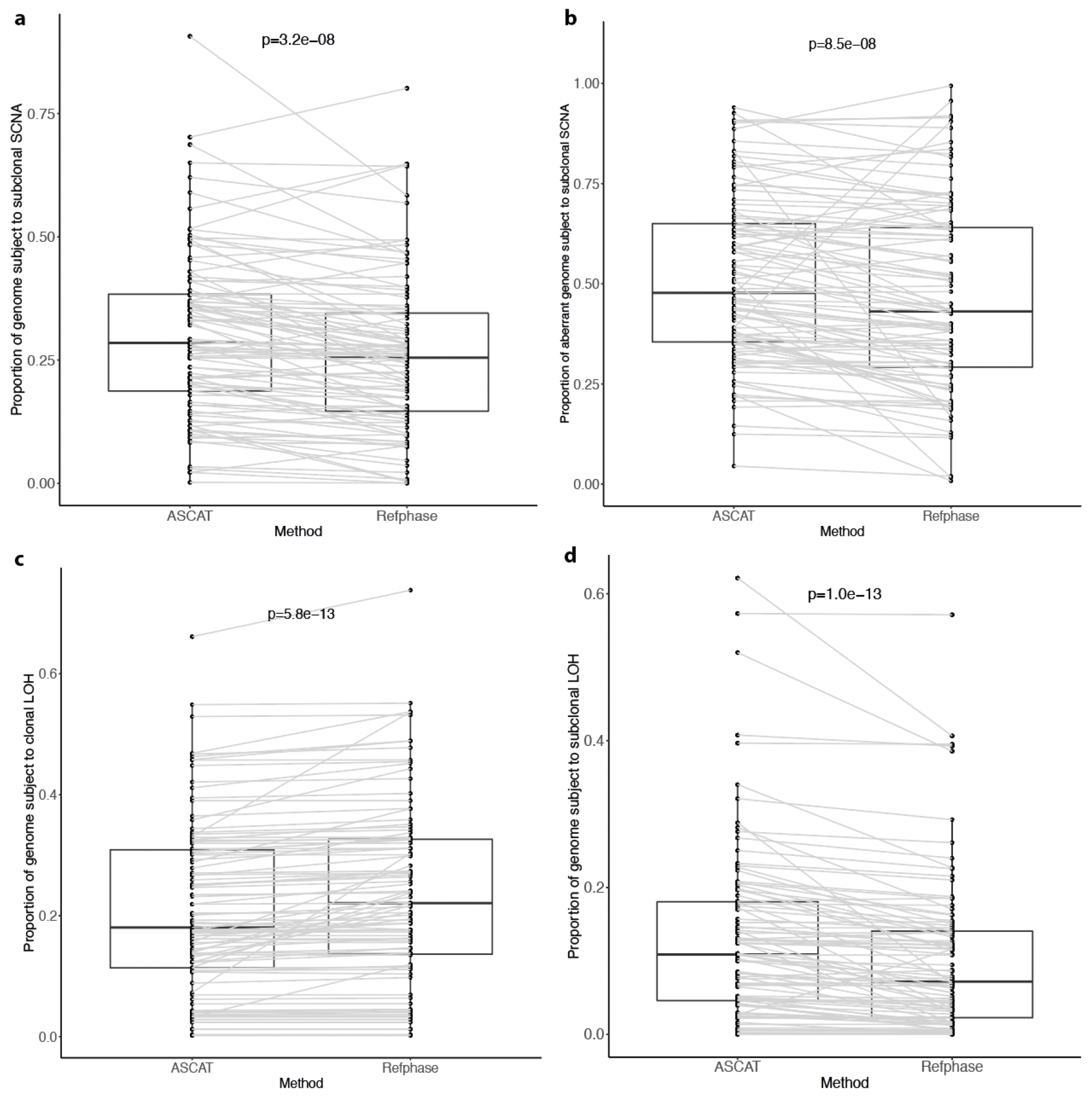
Somatic Copy Number Aberration (SCNA) heterogeneity comparisons between ASCAT and Refphase. **a)** Proportion of the genome subject to subclonal SCNAs (ASCAT median = 0.28, Refphase median = 0.25, p=3.2e-08). **b)** Proportion of aberrant genome subject to subclonal SCNAs, where aberrant genome is defined as the total length of genomic segments in which any SCNA event (clonal or subclonal) is called (ASCAT median = 0.48, Refphase median = 0.43, p=8.5e-08). **c)** Proportion of genome subject to clonal loss of heterozygosity (LOH) (ASCAT median = 0.18, Refphase median = 0.22, p=5.8e-13). **d)** Proportion of the genome subject to subclonal LOH (ASCAT median = 0.11, Refphase median = 0.07, p=1.0e-13). All p-values shown are for paired Wilcoxon signed rank tests with continuity correction, across the n=99 tumours described in Figures 4 and 5. SCNAs in (a) and (b) encompass relative-to-ploidy gains and losses, and LOH events.

**Supplementary Figure 6:**
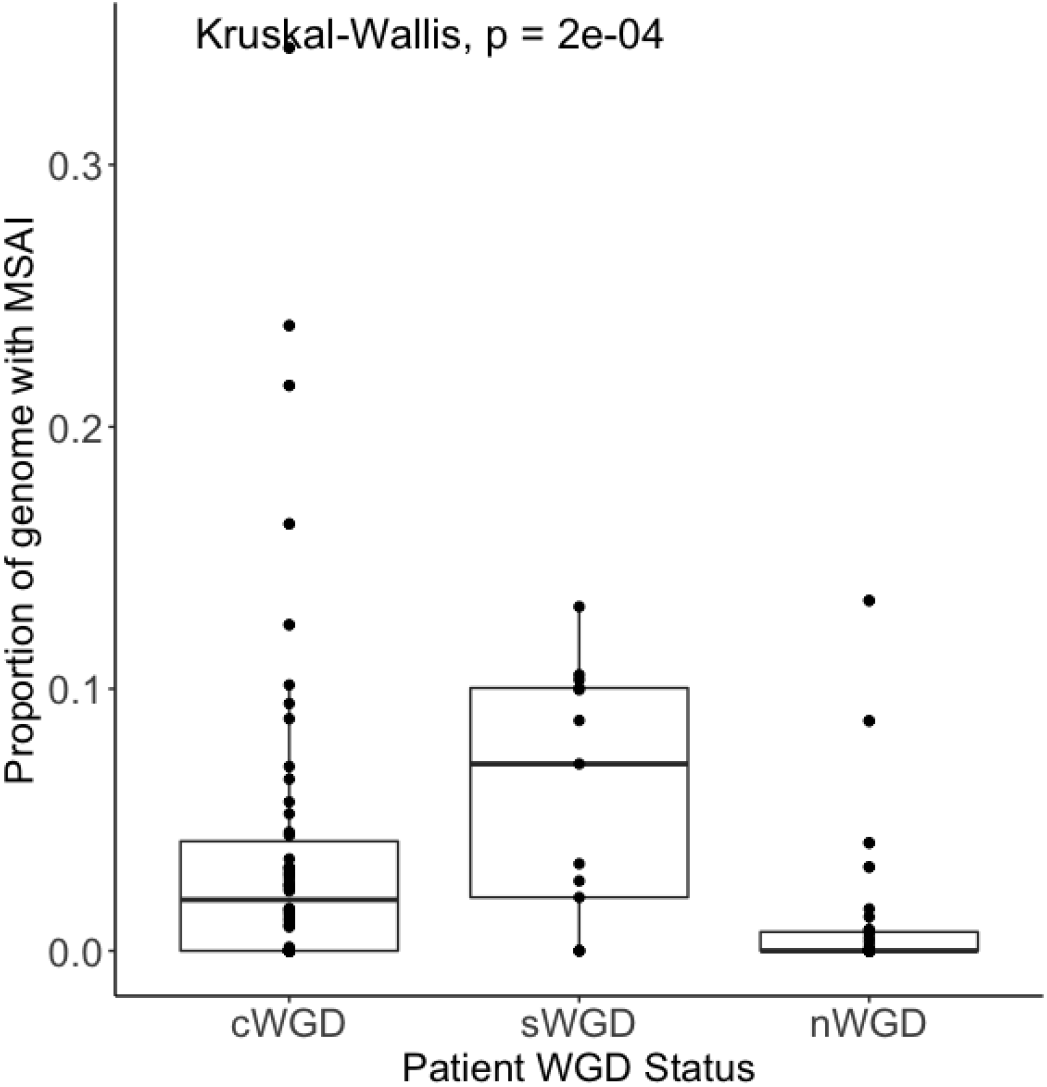
Association between whole genome doubling (WGD) and mirrored subclonal allelic imbalance (MSAI). Median proportions by Patient WGD Status (cWGD = 0.02, n=54 tumours; sWGD = 0.07, n=13 tumours; nWGD = 0, n=32 tumours). Kruskal-Wallis p-value is shown (p=2e-04). Data is shown for n=99 tumours from the pan-cancer cohort showcased in Figure 5 for which MEDICC2 was used to infer WGD status. Proportion of genome data is assessed by Refphase. cWGD = clonal WGD; sWGD = subclonal WGD; nWGD = non-WGD.

**Supplementary Figure 7:**
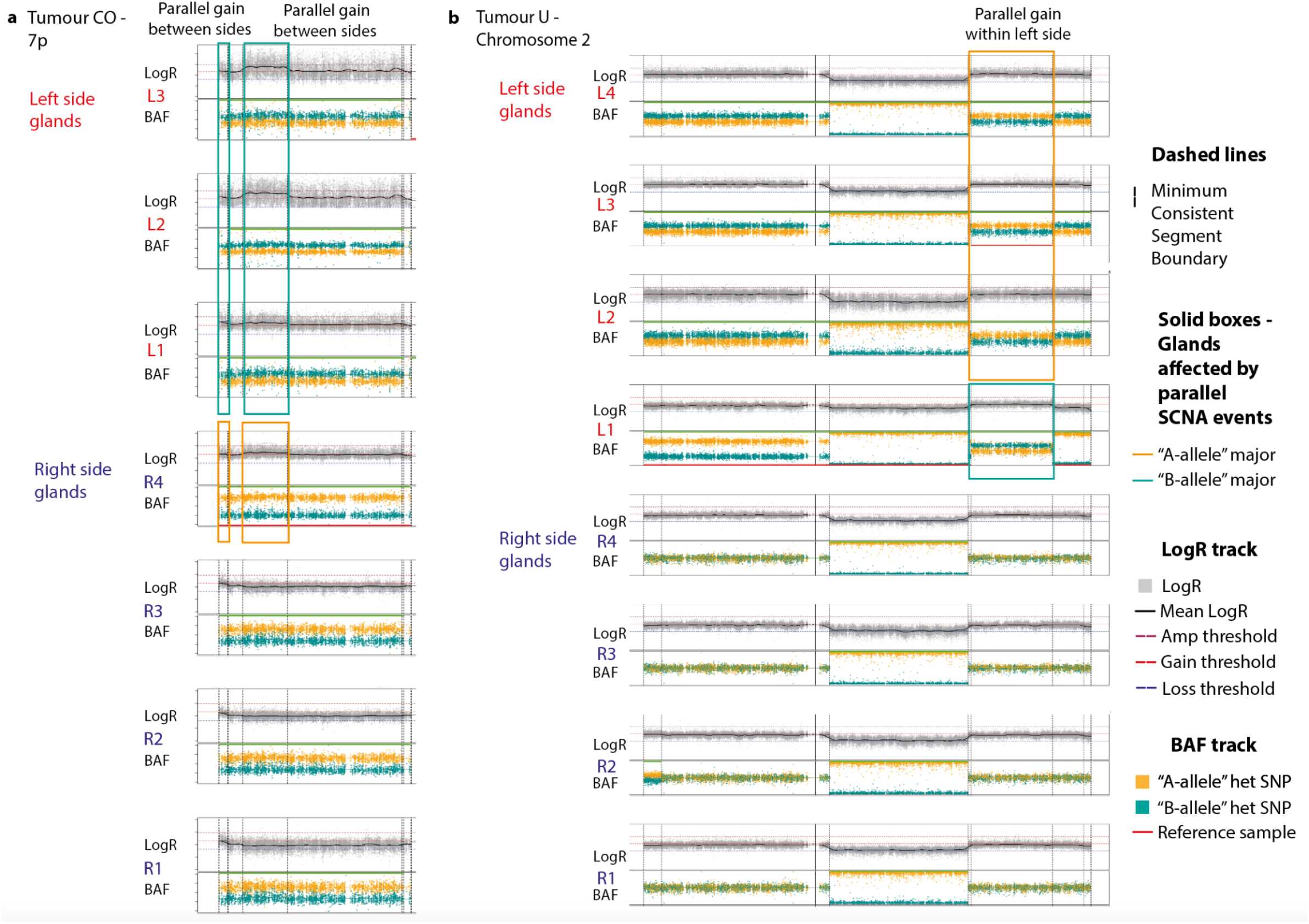
Examples of parallel SCNA events in the Sottoriva *et al* colorectal adenocarcinoma cohort. **a)** Examples of parallel gain events on chromosome arm 7p, observed between glands on different sides of the tumour. **b)** Example of a parallel gain event on chromosome 2, observed within glands on the same side of the tumour. Tumour IDs shown match those in the original publication [44].

**Supplementary Figure 8:**
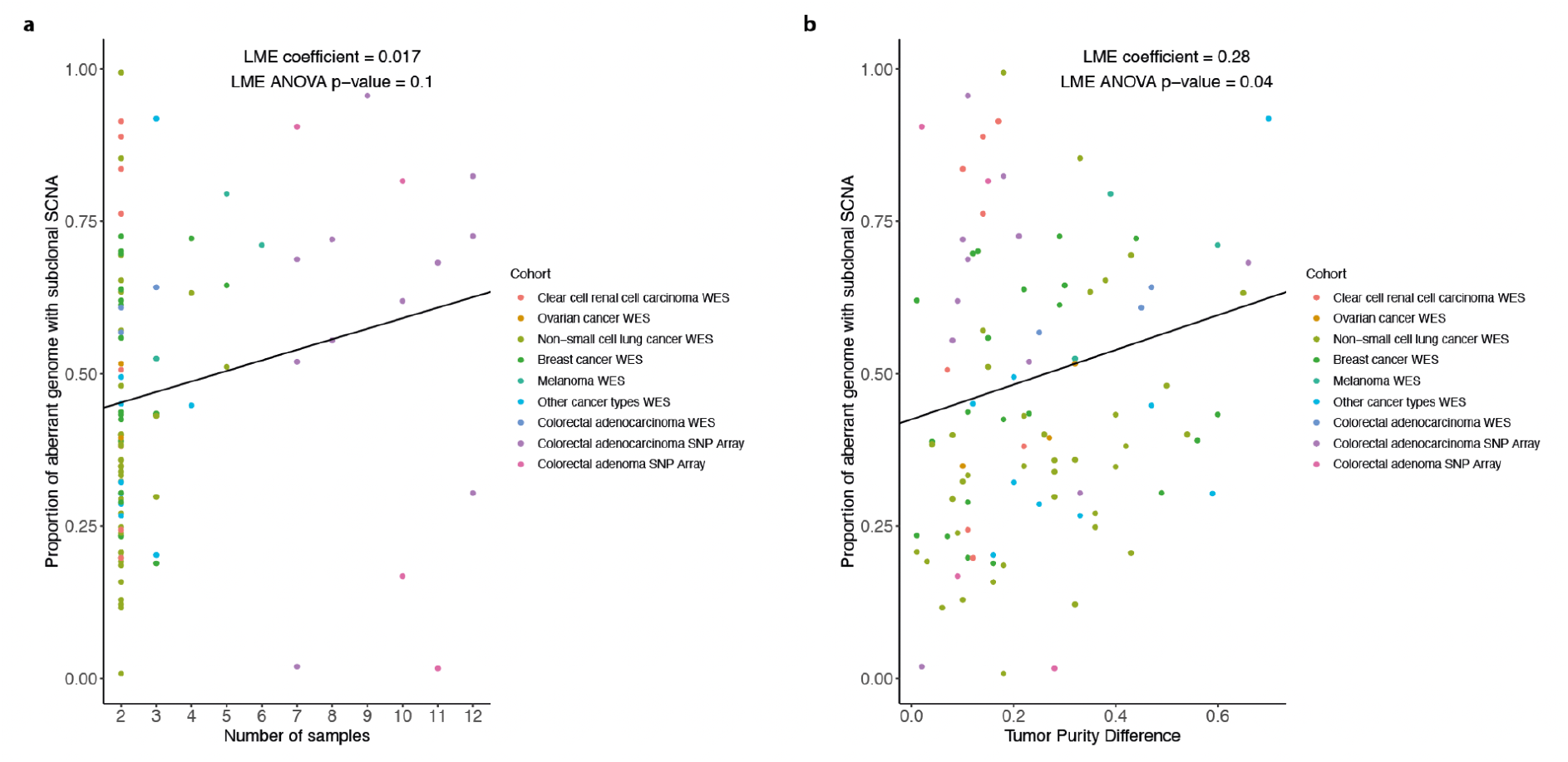
The relationship between SCNA intra-tumour heterogeneity and clinical variables. Association between proportion of aberrant genome with **a)** number of samples (LME coefficient = 0.017, LME ANOVA p=0.1), and **b)** tumour purity difference - defined as the purity difference between the most and least pure sample within a tumour (LME coefficient = 0.28, LME ANOVA p=0.04). Proportion of aberrant genome is defined as the proportion of the total length of genomic segments harbouring any relative-to-ploidy SCNA (gain or loss) or loss of heterozygosity (LOH) event which contains a subclonal SCNA or LOH event. Proportion of the aberrant genome is defined at the tumour level. Analyses are carried out for the pan-cancer cohort described in Figures 4 and 5 (n=99 tumours). Linear mixed effect (LME) ANOVA p-values and LME coefficients calculated using the nlme R package are shown, with analyses adjusted for study cohort (defined by histology and sequencing platform), indicated by the colour legend. Whole-exome sequencing (WES) data is taken from the Brastianos *et al.* study [45]; SNP array data is taken from the Sottoriva *et al.* study [44]. Best fit lines shown have LME slope and intercept values.

